# The molecular mechanism of N-acetylglucosamine side-chain attachment to the Lancefield group A Carbohydrate in *Streptococcus pyogenes*

**DOI:** 10.1101/180810

**Authors:** Jeffrey S. Rush, Rebecca J. Edgar, Pan Deng, Jing Chen, Haining Zhu, Nina M. van Sorge, Andrew J. Morris, Konstantin V. Korotkov, Natalia Korotkova

## Abstract

In many Lactobacillales species (i.e. lactic acid bacteria), peptidoglycan is decorated by polyrhamnose polysaccharides that are critical for cell envelope integrity and cell shape and also represent key antigenic determinants. Despite the biological importance of these polysaccharides, their biosynthetic pathways have received limited attention. The important human pathogen, *Streptococcus pyogenes*, synthesizes a key antigenic surface polymer—the Lancefield group A carbohydrate (GAC). GAC is covalently attached to peptidoglycan and consists of a polyrhamnose polymer, with N-acetylglucosamine (GlcNAc) side chains, which is an essential virulence determinant. The molecular details of the mechanism of polyrhamnose modification with GlcNAc are currently unknown. In this report, using molecular genetics, analytical chemistry and mass spectrometry analysis, we demonstrated that GAC biosynthesis requires two distinct undecaprenol-linked GlcNAc-lipid intermediates: GlcNAc-pyrophosphoryl-undecaprenol (GlcNAc-P-P-Und) produced by the GlcNAc-phosphate transferase GacO and GlcNAc-phosphate-undecaprenol (GlcNAc-P-Und) produced by the glycosyltransferase GacI. Further investigations revealed that the GAC polyrhamnose backbone is assembled on GlcNAc-P-P-Und. Our results also suggested that a GT-C glycosyltranferase, GacL, transfers GlcNAc from GlcNAc-P-Und to polyrhamnose. Moreover, GacJ, a small membrane-associated protein, formed a complex with GacI and significantly stimulated its catalytic activity. Of note, we observed that GacI homologs perform a similar function in *Streptococcus agalactiae* and *Enterococcus faecalis*. In conclusion, the elucidation of GAC biosynthesis in *S. pyogenes* reported here enhances our understanding of how other Gram-positive bacteria produce essential components of their cell wall.

## Introduction

The cytoplasmic membrane of Gram-positive bacteria is surrounded by a thick cell wall consisting of multiple peptidoglycan layers decorated with proteins and a variety of carbohydrate-based polymers. Rhamnose (Rha) is the main component of the carbohydrate structures in many species of the Lactobacillales order (1). These polymers play essential roles in maintaining and protecting bacterial cell envelopes and in pathogenesis of infections caused by *Streptococcus pyogenes* and *Enterococcus faecalis* (1). It is of paramount importance to understand the molecular mechanisms of Rha-containing cell wall polysaccharide biosynthesis for developing novel therapeutics against these bacterial pathogens.

*S. pyogenes* or Group A *Streptococcus* (GAS) is associated with numerous diseases in humans ranging from minor skin and throat infections such as impetigo and pharyngitis to life-threatening invasive infections such as streptococcal toxic syndrome and necrotizing fasciitis (2). The main component of GAS cell wall is the Lancefield group A Carbohydrate (GAC) that comprises about 40-60% of the total cell wall mass (3). Serological grouping of beta-hemolytic streptococci (A, B, C, E, F, and G groups), introduced by Rebecca Lancefield in 1933, is based on the detection of carbohydrate antigens present on the cell wall (4). GAC is presumably covalently linked to N-acetylmuramic acid (MurNAc) of peptidoglycan, although the molecular structure of the linkage unit has not been established (5). All serotypes of GAS produce GAC consisting of a polyrhamnose backbone with N-acetylglucosamine (GlcNAc) side chains (6,7). The Lancefield classification of GAS relies on serological specificity of the immunodominant GlcNAc residues in the carbohydrate antigen (7). The average molecular mass of GAC has been reported to be 8.9±1.0 kDa, corresponding to an average of 18 repeating units of [→3)αRha(1→2)[β-GlcNAc(1→3)]α-Rha(1→] (8). Since Rha is absent in mammalian cells, GAC is an attractive candidate for a universal GAS vaccine (8,9). Moreover, the GlcNAc side chains are an important virulence determinant in GAS (5). GAS mutants lacking GlcNAc are susceptible to innate immune clearance by neutrophils and antimicrobial agents, and are significantly attenuated in animal models of GAS infection (5).

Similar to the biosynthesis of other cell-envelope polymers — peptidoglycan, lipopolysaccharide, wall teichoic acid (WTA), and capsular polysaccharide — the synthesis of Rha-containing carbohydrate structures is likely initiated on the inside of the cytoplasmic membrane and proceeds through several steps including the attachment of the first sugar residue to a lipid carrier, undecaprenyl phosphate (Und-P), followed by elongation of the polysaccharide through stepwise addition of activated sugar residues to the lipid carrier (1). After translocation of the polysaccharide across the membrane and further modifications, it is presumably attached to peptidoglycan via a phosphate ester linkage (1). The first membrane step of O-antigen biosynthesis in enterobacteria and WTA biosynthesis in *Bacillus subtilis* and *Staphylococcus aureus* (10) is catalyzed by the UDP-GlcNAc:Und-P GlcNAc-1-phosphate transferase encoded by WecA homologs (11,12). WecA transfers GlcNAc-phosphate from UDPGlcNAc to Und-P, forming GlcNAcpyrophosphoryl-undecaprenol (GlcNAc-P-P-Und). Although GAS strains do not produce WTA, all GAS genomes contain a *wecA* homolog, *gacO* (5). Significantly, GAC biosynthesis is sensitive to tunicamycin, a known inhibitor of WecA (5) and the *Streptococcus mutans* GacO homolog, RgpG, has been shown to complement WecA activity in *E. coli* (13). These observations support an essential role for GacO in the biosynthesis of GlcNAc-P-P-Und and suggest GlcNAc-P-P-Und may function as a lipid-anchor in the initiation of GAC biosynthesis.

Other genes required for GAC biosynthesis and transport are found in a separate location on the GAS chromosome and comprise a 12-gene locus (*gacA-gacL*) (Fig.1A). The first three genes, *gacA*-*gacC* together with *gacG,* are conserved in many species of the Lactobacillales order (1). *GacA* encodes a dTDP-4-dehydrorhamnose reductase, the enzyme responsible for dTDP-rhamnose biosynthesis (14). *GacB*, *gacC*, and *gacG* encode putative cytoplasmic rhamnosyltransferases. In *S. mutans* homologous genes *rgpA*, *rgpB* and *rgpF* are involved in rhamnan backbone biosynthesis (15). In GAS, *gacA*-*gacC* are essential for viability and cannot be deleted (5). However, deletion mutants have been obtained in other genes of the GAC gene cluster (5). It has been shown that *gacI*, *gacJ* and *gacK* are non-essential for viability, but are required for GlcNAc side-chain addition to the polyrhamnose (5). They encode a putative cytoplasmic glycosyltransferase, a small membrane protein and a protein with homology to the Wzx family of membrane proteins involved in the export of O-antigen and teichoic acids, respectively. In contrast, inactivation of *gacD*, *gacE*, *gacF*, *gacG*, *gacH* or *gacL* has no effect on GAS viability and the GAC produced by these mutants is reported to display a wild type (WT) antigenic profile, indicating the presence of the immunodominant GlcNAc side chains (5). *GacD* and *gacE* encode the components of an ABC transport system. In *S. mutans* the *gacD* and *gacE* homologs are responsible for rhamnan polysaccharide transport (15). *GacF*, *gacH* and *gacL* encode a cytosolic glycosyltransferase, a putative membrane-associated glycerol phosphate transferase and a membrane-associated glycosyltransferase, respectively.

**Figure 1.**
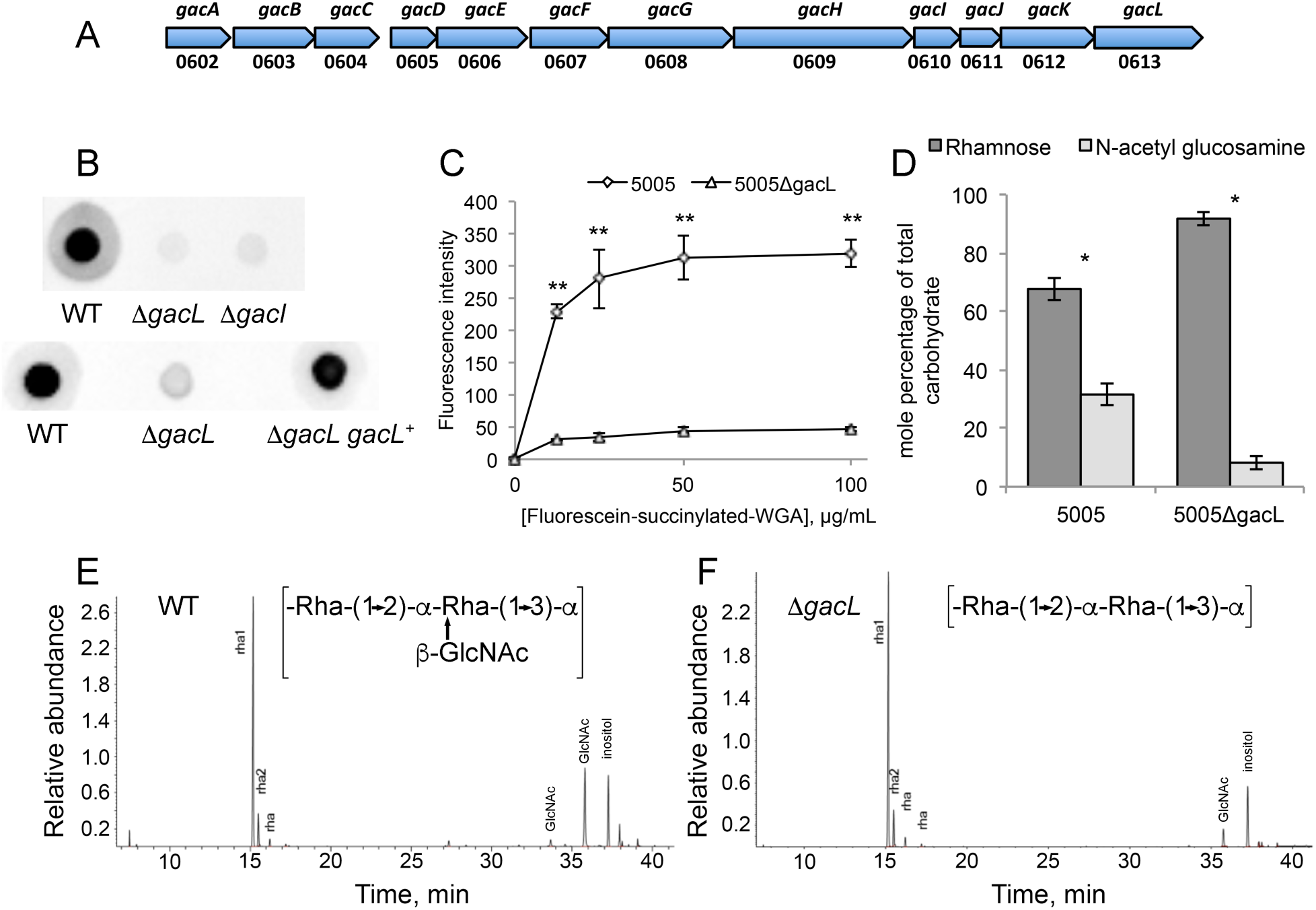
Map of *S. pyogenes* genes involved in Group A Carbohydrate (GAC) biosynthesis and analysis of 5005Δ*gacI* and 5005Δ*gacL* deletion mutants. (A) GAC biosynthesis gene cluster. Numbers below represent MGAS5005 gene designations (B) Representative immunoblot analysis of cell-wall fractions isolated from MGAS5005, 5005Δ*gacI*, 5005Δ*gacL* and 5005Δ*gacL gacL*^*+*^. Data are representative of biological triplicates. (C) Binding of N-acetyl glucosamine specific fluorescein-succinylated-WGA to whole MGAS5005 and 5005Δ*gacL* was measured. Data are the average of three replicates ± standard deviation. (D) Rhamnose and GlcNAc mole percentage of total carbohydrate was determined by GC-MS for cell wall material isolated from MGAS5005 and 5005Δ*gacL* following methanolysis as described in Experimental procedures. Data are the average of four replicates ± standard deviation. The asterisk indicate statistically different values (* p < 0.05, ** p < 0.01) as determined by the Student’s t-test. (E and F) GC-MS chromatograms for glycosyl composition analysis of cell wall isolated MGAS5005 and 5005Δ*gacL*. The deduced schematic structure of the repeating unit of GAC is shown for each strain. The chromatograms are representative of four separate analyses performed on two different cell wall preparations.

In this study we address the molecular mechanism of GlcNAc attachment to polyrhamnose. We demonstrate that GacI catalyzes formation of GlcNAc-P-Und, and GacJ stimulates the catalytic activity of GacI. Subsequently GacL transfers GlcNAc from GlcNAc-P-Und to the polyrhamnose backbone of GAS polysaccharide. Moreover, we confirm GacO function in GlcNAc-P-P-Und formation and demonstrate the role of this GlcNAc-lipid in initiation of polyrhamnose biosynthesis.

## Results

### GacL is required for GlcNAc attachment to polyrhamnose

GacL, a polytopic (12–13 transmembrane segments) membrane protein, is reported to be dispensable for GlcNAc attachment to polyrhamnose (5). To investigate the function of GacL in GAC biogenesis, we disrupted *gacL* in the hyperinvasive *S. pyogenes* M1T1 serotype strain, MGAS5005 (16), creating the 5005Δ*gacL* mutant. In agreement with published data (5) 5005Δ*gacL* did not display any detectable growth phenotype in comparison to the WT strain. Surprisingly, we found that 5005Δ*gacL* cells failed to bind GlcNAc-specific anti-GAC antibodies, suggesting a loss of the GlcNAc antigenic epitope (Fig. 1B). The 5005Δ*gacL* phenotype was restored by expressing the WT copy of *gacL* on the mutant chromosome (Fig. 1B). To confirm that 5005Δ*gacL* cells are deficient in GlcNAc addition to polysaccharide, we measured the binding of fluorescently labeled succinylated wheat germ agglutinin (sWGA), a lectin that specifically binds non-reducing terminal β-GlcNAc residues, to WT and 5005Δ*gacL* cells. Deletion of *gacL* led to a significant decrease in binding of WGA to the 5005Δ*gacL* strain as compared with the WT (Fig. 1C), indicating that the mutant has substantially less GlcNAc-containing saccharides on the cell surface. Furthermore, direct compositional analysis of cell wall purified from WT and 5005Δ*gacL* cells confirmed a significant decrease of GlcNAc content in the 5005Δ*gacL* cell wall sample (Fig.1D, E and F). These data strongly support a role for GacL in GlcNAc side-chain attachment to polyrhamnose.

### A *gacL* deletion mutant accumulates GlcNAc-P-Und

To investigate further the effect of GacL deletion in *S. pyogenes*, we isolated the phospholipid fractions from the membranes of WT and 5005Δ*gacL* cells. TLC analysis revealed accumulation of a previously unidentified phosphoglycolipid in 5005Δ*gacL* (Fig. 2A). The novel lipid was found to be stable to mild alkaline methanolysis, but sensitive to mild acid (0.1 N HCl, 50 °C, 50% isopropanol) and reacted with orcinol spray (17), Dittmer-Lester phospholipid spray reagent (18) and with anisaldehyde, an isoprenol-specific reagent (19), consistent with its tentative identification as a glycophosphoprenol (data not shown). High-resolution negative ion ESI-MS analysis of the purified lipid identified a molecular ion [M-H]^-1^ of M/z = 1048.73621 (Fig. 2B). In addition, prominent high-resolution fragment ions characteristic of PO_3_, H_2_PO_4_, N-acetyl hexosamine-phosphate and Und-P were also found (Table 1). These data are consistent with a glycolipid comprised of N-acetyl hexosamine-phosphate-undecaprenol. In experiments described in detail below, we found that when membrane fractions from GAS are incubated with UDP-[^3^H]GlcNAc, a [^3^H]GlcNAc-lipid which co-chromatographs on TLC with this novel lipid is rapidly formed. Importantly, the product ion spectra shown in Figure 2B does not contain a fragment ion containing [Und-PO_4-_C_2_H_2_NHCOCH_3_]^-^ (M/z=929.7), arising from a cross-ring fragmentation reaction, suggesting that the hydroxyl of the anomeric carbon and the nitrogen at the 2-position of the glycosyl ring are *trans* to the plane of the ring (20,21). This observation strongly suggests that the anomeric hydroxyl is present in the β configuration. Furthermore, isolated [^3^H]GlcNAc-P-Und phospholipid (described in detail below) is extremely sensitive to incubation with 50% phenol at 68 °C — a property that is consistent with β-GlcNAc-P-Und (22) (Table S1). Taken together, these data identify the novel glycolipid accumulating in the 5005Δ*gacL* strain as β-GlcNAc-P-Und and suggest that GacL is a GlcNAc transferase using β-GlcNAc-P-Und as GlcNAc donor for the addition of the GlcNAc side-chains to GAC.

**Figure 2.**
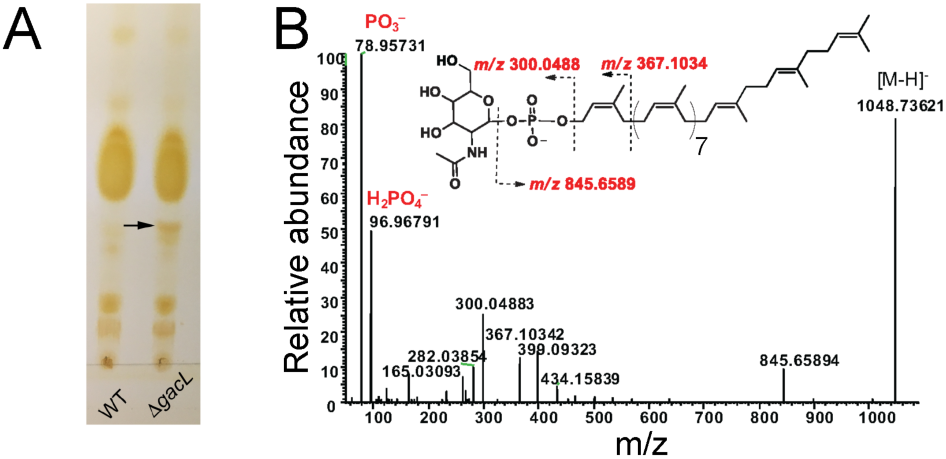
Purification and identification of GlcNAc-phosphate-undecaprenol in 5005Δ*gacL.* (A) Thin layer chromatography (TLC) analysis of phospholipids isolated from MGAS5005 and 5005Δ*gacL*. Phospholipids extracted from bacterial strains were separated by TLC on Silica Gel G in CHCl_3_/CH_3_OH/H_2_O/NH_4_OH (65:25:4:1). Position of the novel, alkaline-resistant and acid-labile, phospholipid accumulating in 5005Δ*gacL* is indicated by the arrow. The results are representative of three separate experiments. (B) ESI-MS/MS analysis of the novel phospholipid isolated from 5005Δ*gacL*. The spectrum is assigned to GlcNAc-phosphate-undecaprenol.

**Table 1.** The major fragments of the precursor ion at *m/z* 1048.

| $m/z$ | Formula | Mass error (ppm) |
| --- | --- | --- |
| 78.95731 | PO<sub>3</sub> | -8.19 |
| 96.96791 | H<sub>2</sub>PO<sub>4</sub> | -6.31 |
| 300.04883 | C<sub>8</sub>H<sub>15</sub>O<sub>9</sub>NP | 3.12 |
| 845.65894 | C<sub>58</sub>H<sub>88</sub>ONP | -1.02 |

Fragment ions originating from the precursor molecular ion of 1048 were subjected to MS/MS fragmentation as described in Experimental procedures. The proposed molecular formulae of the product ions are supported by the observed isotope patterns for the relevant ions.

### GAS synthesizes two GlcNAc-lipids *in vitro*

To investigate the biosynthetic origin of the novel glycolipid accumulating in 5005Δ*gacL* cells, membrane fractions from MGAS5005 were incubated with UDP-[^3^H]GlcNAc and analyzed for lipid products as described in Experimental procedures. Experiments showed that [^3^H]GlcNAc was efficiently transferred from UDP-[^3^H]GlcNAc into two detectable GlcNAc-lipids (Fig. 3A, Table 2). The major lipid product co-migrates on TLC with the novel lipid accumulating in the 5005Δ*gacL* strain and is identified as β-[^3^H]GlcNAc-P-Und, as described above. The minor product is assumed to be [^3^H]GlcNAc-P-P-Und, since it co-migrates with authentic GlcNAc-P-P-Und synthesized in *B. cereus* membranes (Fig. 3B) (23) and its formation is potently inhibited by tunicamycin (Fig. S1).

**Figure 3.**
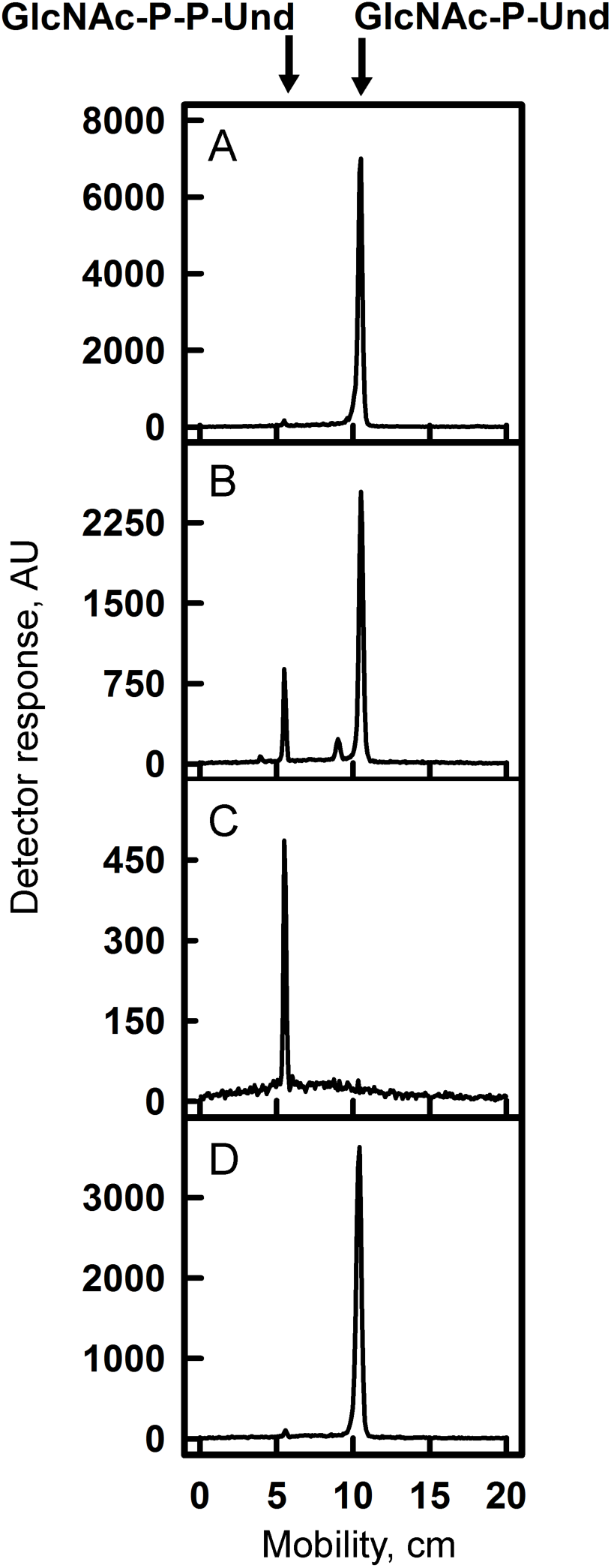
Thin layer chromatography of [^3^H]GlcNAc-lipids from *in vitro* incubations of GAS mutants and *Bacillus cereus* membranes with UDP-[^3^H]GlcNAc. Membrane fractions from MGAS5005 (panel A), *B. cereus* (Panel B), 5005Δ*gacI* (Panel C), or 5005Δ*gacL* (Panel D) were incubated with UDP-[^3^H]GlcNAc and analyzed for [^3^H]GlcNAc lipid synthesis by thin layer chromatography. Reaction mixtures contained 50 mM Tris-Cl, pH 7.4, 5 mM 2-mercaptoethanol, 20 mM MgCl_2_, 1 mM ATP, 5 μM UDP-[^3^H]GlcNAc (486 cpm/pmol) and bacterial membrane suspension (100-200 μg membrane protein) in a total volume of 0.02 ml. Following a 10 minute pre-incubation at 30°C, GlcNAc-lipid synthesis was initiated by the addition of UDP-[^3^H]GlcNAc. After 10 min, reactions were processed for GlcNAc-lipid synthesis as described in Experimental procedures. The organic layers were dried, dissolved in a small volume of CHCl_3_/CH_3_OH (2:1) and a portion was removed and assayed for radioactivity by liquid scintillation spectrometry. The remainder was spotted on 10 x 20 cm plate of silica gel G and developed in CHCl_3_/CH_3_OH/NH_4_OH/H_2_O (65:25:1:4). [^3^H]GlcNAc-lipids were detected by scanning with an AR2000 Bioscan Radiochromatoscanner. AU, arbitrary units. The results are representative of three separate experiments.

**Table 2.** GlcNAc-lipid synthesis in membrane fractions from various bacterial strains.^a^

| Enzyme Source | GlcNAc-P-Und,<br>pmol/mg | GlcNAc-P-P-Und,<br>pmol/mg |
| --- | --- | --- |
| MGAS5005 | 259.4±8.2 | 7.8±1.8 |
| 5005Δ*gacI* | Not Detected | 15.9±2.5 |
| GBS COH1 | 62±0.4 | 5.6±0.8 |
| GBSΔ*gacI* | Not Detected | 12±2 |
| *E. coli* Rosetta<br>(DE3) | Not Detected | 19.5±0.3 |
| *E. coli*:GacI | 102.8±3.3 | 5.4±0.05 |
| *E. coli*:EpaI | 189±3.9 | 7±1.5 |

<sup>a</sup>Reaction mixtures contained 50 mM Tris-HCl, pH 7.4, 5 mM 2-mercaptoethanol, 20 mM MgCl<sub>2</sub>, 1 mM ATP, 5 μM UDP-[<sup>3</sup>H]GlcNAc (486 cpm/pmol) and bacterial membrane suspension (50-100 μg membrane protein) in a total volume of 0.01 ml. Following a 10 minute pre-incubation at 30°C, GlcNAc-lipid synthesis was initiated by the addition of UDP-[<sup>3</sup>H]GlcNAc. After 5 min, reactions were processed for GlcNAc-lipid synthesis as described in Experimental procedures. The results are representative of at least three separate experiments.

### GacI encodes a UDP-GlcNAc:Und-P GlcNAc transferase activity

The GAS polysaccharide gene cluster contains a gene*, gacI*, reported to be required for GlcNAc addition to cell wall polysaccharide (5) and annotated as a GT-A type glycosyltransferase. The following studies were conducted to determine if GAS GacI might be responsible for the synthesis of GlcNAc-P-Und. To determine if GacI function was essential for formation of GlcNAc-P-Und, we generated a deletion of *gacI* in the MGAS5005. In agreement with published data (5) 5005Δ*gacI* lost reactivity with anti-GAC antibodies, with specificity for the immunodominant GlcNAc, indicating a loss of GlcNAc modification in polyrhamnose (Fig. 1B). When 5005Δ*gacI* membrane fractions were incubated with [^3^H]UDP-GlcNAc no incorporation into [^3^H]GlcNAc-P-Und was observed and only the minor lipid, [^3^H]GlcNAc-P-P-Und was found (Fig. 3C, Table 2), strongly supporting an essential role for GacI in GlcNAc-P-Und synthesis. The observation that the 5005Δ*gacL* strain synthesizes normal levels of [^3^H]GlcNAc-P-Und *in vitro* (Fig. 3D) indicates that 5005Δ*gacL* does not fail to add GlcNAc side-chains to polyrhamnose due to a lack of GlcNA-P-Und and supports the conclusion that GacL may be the GlcNAc-P-Und:polyrhamnan GlcNAc transferase.

To confirm the role of GacI in GlcNAc-P-Und biosynthesis we expressed GAS GacI in *E. coli*. When membrane fractions from *E. coli* carrying an empty vector were incubated with UDP-[^3^H]GlcNAc and analyzed by silica gel TLC, a small peak of [^3^H]GlcNAc-P-P-Und was found (Table 2, Fig. S2A). In contrast, the membranes of recombinant *E. coli* expressing GacI accumulated two products corresponding to a small amount of GlcNAc-P-P-Und and a very large amount of GlcNAc-P-Und (Table 2, Fig. S2B). Thus, our data strongly indicate that GacI is the GlcNAc-P-Und synthase.

### GacI homologs in GBS and *E. faecalis* function in the biosynthesis of GlcNAc-P-Und

The presence of GlcNAc-P-P-Und and GlcNAc-P-Und have been previously reported in *B. cereus* and *Bacillus megaterium* (12,23). Our *in vitro* studies show that the MGAS5005 strain synthesizes two [^3^H]GlcNAc-lipids that exactly co-migrate on TLC with the previously reported [^3^H]GlcNAc-lipids from *B. cereus* (Fig. 3A and 3B). Moreover, an analysis of bacterial genomes using GacI in a BLAST search identified a GacI homolog (60% identity) in *B. cereus* and *B. megaterium*, suggesting its role in GlcNAc-P-Und biosynthesis in these bacteria (supplemental file 1). Furthermore, *gacI* homologs were identified in the polysaccharide biosynthesis gene clusters of the important human pathogens *E. faecalis* (*epaI* with 48% sequence identity) (24) and *S. agalactiae* (Group B Streptococcus or GBS) (SAN_1536 gene with 46% sequence identity) (Fig. S3). To investigate whether the GBS homolog of GacI also catalyzes the formation of GlcNAc-P-Und, we engineered a knock out of the GacI homolog in GBS COH1. Although GBS COH1 membranes synthesize both [^3^H]GlcNAc-P-P-Und and [^3^H]GlcNAc-P-Und (Fig. 4B and Fig. S2D), GBS COH1Δ*gacI* membranes no longer synthesize [^3^H]GlcNAc-P-Und (Table 2 and Fig. S2E).

**Figure 4.**
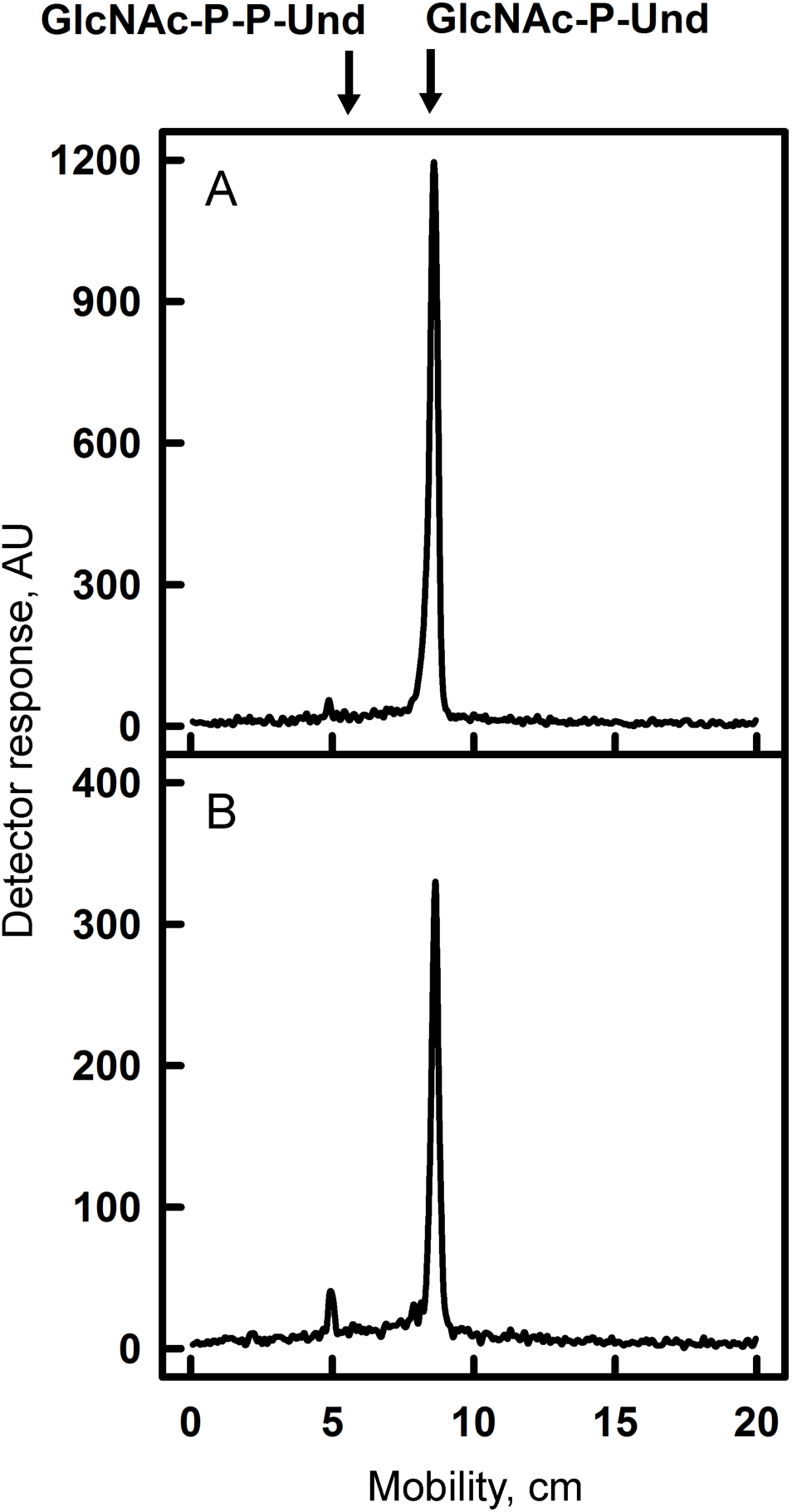
Thin layer chromatography of [^3^H]GlcNAc-lipids from *in vitro* incubations of GAS and GBS membranes with UDP-[^3^H]GlcNAc. Membrane fractions from WT MGAS5005 (panel A) or GBS COH1 (Panel B) were incubated with UDP-[^3^H]GlcNAc and analyzed for [^3^H]GlcNAc-lipid synthesis by thin layer chromatography. Reaction mixtures were exactly as described in the legend to Figure 3. After 5 min incubation with UDP-[^3^H]GlcNAc, reactions were processed for GlcNAc-lipid synthesis as described in Experimental procedures. The organic layers were dried, dissolved in a small volume of CHCl_3_/CH_3_OH (2:1) and a portion was removed and assayed for radioactivity by liquid scintillation spectrometry. The remainder was spotted on 10 x 20 cm plate of silica gel G and developed in CHCl_3_/CH_3_OH/NH_4_OH/H_2_O (65:25:1:4). [^3^H]GlcNAc-lipids were detected by scanning with an AR2000 Bioscan Radiochromatoscanner. AU, arbitrary units. The results are representative of three separate experiments.

To confirm that the GacI homolog detected in *E. faecalis*, EpaI (24), also possesses GlcNAc-P-Und synthase activity, EpaI was expressed exogenously in *E. coli* and found to actively catalyze the formation of [^3^H]GlcNAc-P-Und (Table 2, Fig. S2C). Altogether, our results are consistent with the function of GacI homologs from GAS, GBS and *E. faecalis* in the transfer of GlcNAc from UDP-GlcNAc to Und-P forming GlcNAc-P-Und.

### ATP stimulates GlcNAc-P-Und biosynthesis

Preliminary enzymatic properties for the synthesis of [^3^H]GlcNAc-P-Und established that Mg^2+^ was the preferred divalent cation and that the formation of GlcNAc-P-Und was substantially stimulated by exogenously added Und-P (as a dispersion in CHAPS detergent). In early studies of GAC biosynthesis in GAS (25), it was shown that the addition of ATP dramatically stimulated incorporation of radioactive UDP-[^3^H]GlcNAc into [^3^H]GlcNAc membrane lipids and [^3^H]GlcNAc-polysaccharide. We confirmed that inclusion of 1 mM ATP significantly stimulated the incorporation of [^3^H]GlcNAc into [^3^H]GlcNAc-P-Und (Fig. 5 and Fig. S4), as well as [^3^H]GlcNAc-P-P-Und (Fig. S4). However, the inclusion of ATP did not stimulate GacI activity in *in vitro* assays of CHAPS-soluble, affinity purified GacI solely dependent on exogenously added Und-P as acceptor. These data suggest that the effect of ATP addition is most likely due to formation of Und-P, *in situ,* by phosphorylation of endogenous undecaprenol by undecaprenol kinase (26).

**Figure 5.**
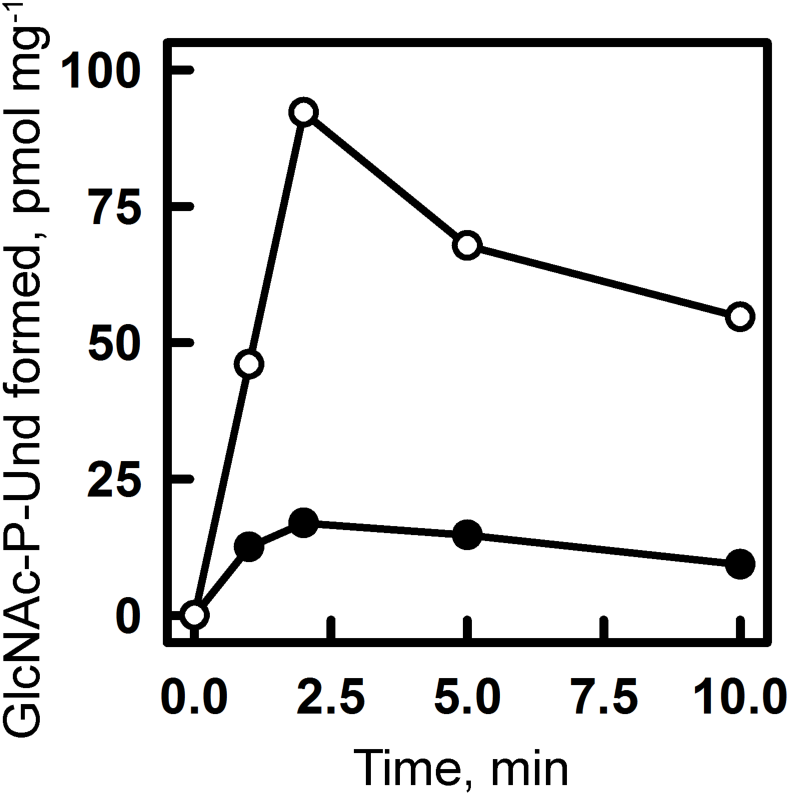
Effect of ATP on GlcNAc-P-Und synthesis *in vitro* in MGAS5005 membrane fractions. GlcNAc-P-Und synthesis in MGAS5005 membranes was assayed after the indicated time at 30 °C in the presence (∘) or absence (•) of 1 mM ATP. Reaction mixtures were identical to those described in Figure 3 except for the presence of ATP. Following incubation, incorporation into [^3^H]GlcNAc-P-Und was determined as described in Experimental procedures. The results are representative of three separate experiments.

### GacJ forms a complex with GacI and enhances its catalytic efficiency

*GacJ* is located immediately downstream of *gacI* in the GAS GAC biosynthesis gene cluster and encodes a small, 113 amino acid, membrane protein (Fig. 1A). We hypothesized that GacI might form an obligate complex with GacJ. This hypothesis is based on the observation that *gacJ* and *gacI* are frequently located adjacent to each other on bacterial chromosomes and are sometimes fused to form a single polypeptide (accession numbers: GAM11018, ADH85075, ALC15489 and ADU67183). To test this hypothesis, we solubilized membranes from *E. coli* co-expressing GacJ and amino-terminal His-tagged GacI with the zwitterionic detergent CHAPS, and isolated GacI complexes using Ni-NTA chromatography (Fig. 6A). SDS-PAGE of the affinity-purified sample revealed two bands corresponding to the anticipated molecular sizes of GacI and GacJ (Fig. 6B). Proteomics analysis of the excised protein bands confirmed the identities of the recovered proteins. This result indicates that GacI and GacJ form a stable, CHAPS-soluble complex and co-purify during affinity chromatography.

**Figure 6.**
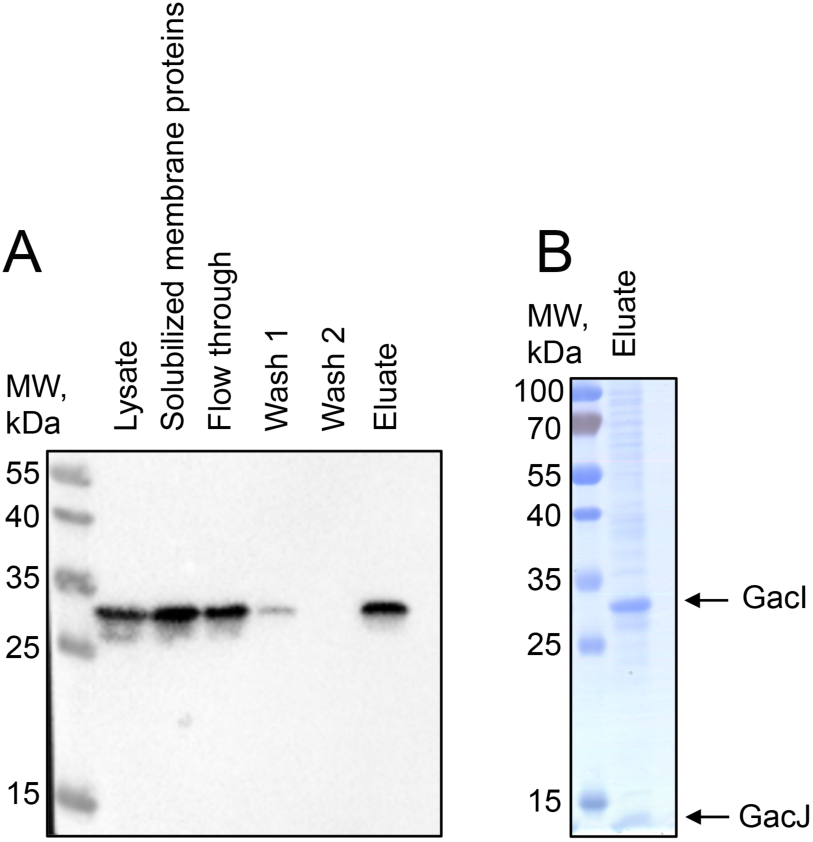
GacI and GacJ exist as a detergent-stable complex in the membrane. GacJ and His-tagged GacI were co-expressed in *E. coli* Rosetta DE3 cells and extracted from the membrane fraction in 2.5% CHAPS. The proteins were purified using Ni-NTA agarose in the presence of 2.5% CHAPS. (A) Fractions collected during Ni-NTA purification were analyzed by immunoblot using anti-His antibodies. (B) The eluted proteins were analyzed by SDS-PAGE. The results are representative of three separate experiments.

To investigate if GacJ performs any catalytic function in association with GacI, we tested the proteins for GlcNAc-P-Und synthase activity *in vitro*, individually and in combination. When GacI was expressed singly in *E. coli*, enzymatically active protein was found in the membrane fraction (Table 2, Fig. S2B), indicating that GacI does not require GacJ for activity or membrane association. GacJ was found to be catalytically inactive when expressed by itself in *E. coli*, consistent with the observation that 5005Δ*gacI* membranes show no residual GlcNAc-P-Und synthase activity. Kinetic analysis of GlcNAc-P-Und synthase activity for Und-P in membrane fractions of *E. coli* expressing GacI revealed an apparent Km of 18.7 μM (Table 3, Fig. S5A). However, co-expression of GacJ dramatically lowered the apparent Km of GacI for Und-P to 1.1 μM, suggesting a significant change in affinity for the lipid acceptor. Moreover, the V_max_ of GacI activity increased from 54.2 pmol/min/mg to 18.5 nmol/min/mg in the presence of GacJ (Table 3, Fig. S5B). A kinetic analysis of GlcNAc-P-Und synthase for Und-P in MGAS5005 strain gave an apparent Km of 6.4 μM and a V_max_ of 333 pmol/min/mg (Table 3, Fig. S5C), similar to the enzymatic parameters of GacI co-expressed with GacJ in *E. coli*. These enzymatic parameters are in marked contrast to those of the GlcNAc-P transferase that synthesizes GlcNAc-P-P-Und. When a similar kinetic analysis was performed in 5005Δ*gacI*, so that only GlcNAc-P-P-Und synthesis could be scored, an apparent Km for Und-P of 19.3 μM was obtained and the V_max_ was only 2.75 pmol/min/mg (Table 3, Fig. S5D). Clearly, GacI displays a much greater catalytic efficiency for the synthesis of GlcNAc-P-Und than the enzyme synthesizing GlcNAc-P-P-Und, which is reflected in the dramatic difference in the rates of synthesis of the two glycolipids. In summary,our data indicate that GacJ forms a stable association with GacI and stimulates the catalytic activity of GacI.

**Table 3.**
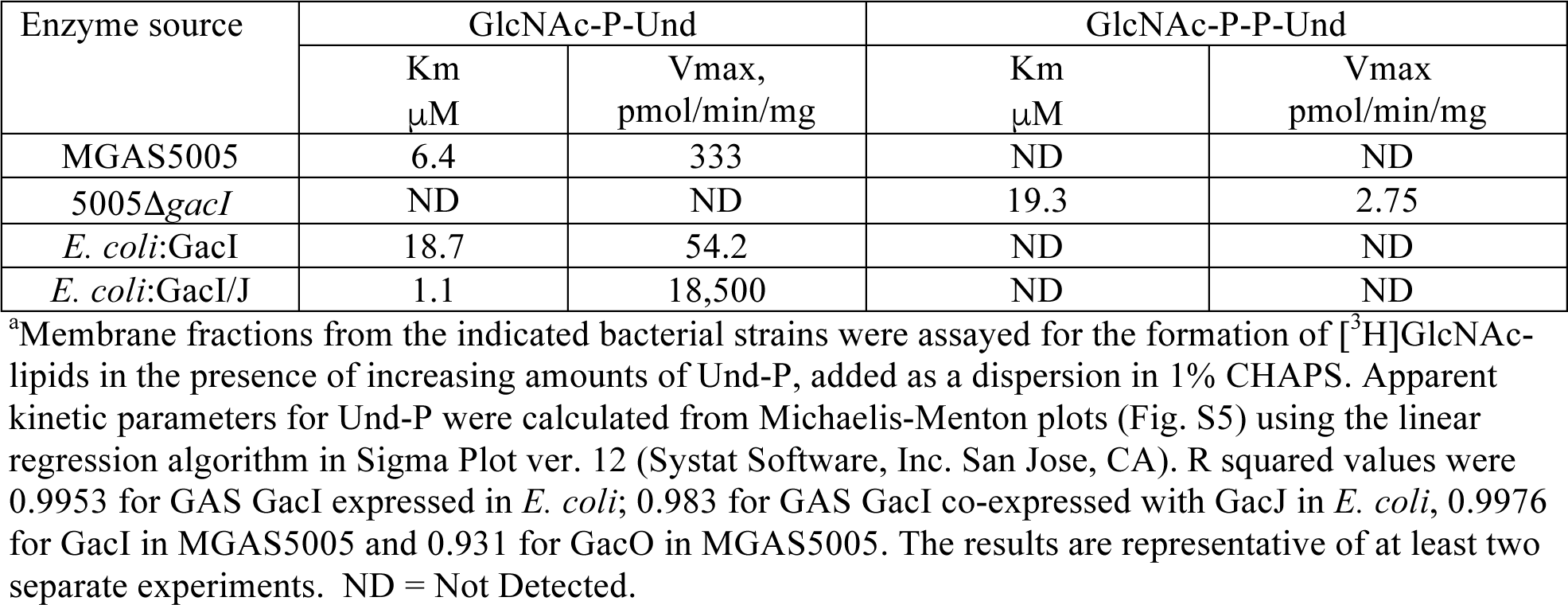
Kinetic parameters of GlcNAc-lipid synthases in membrane fractions from various bacterial strains.^a^

| Enzyme source | GlcNAc-P-Und |  | GlcNAc-P-P-Und |  |
| --- | --- | --- | --- | --- |
|  | Km<br>μM | Vmax,<br>pmol/min/mg | Km<br>μM | Vmax<br>pmol/min/mg |
| MGAS5005 | 6.4 | 333 | ND | ND |
| 5005Δ*gacI* | ND | ND | 19.3 | 2.75 |
| *E. coli*:GacI | 18.7 | 54.2 | ND | ND |
| *E. coli*:GacI/J | 1.1 | 18,500 | ND | ND |

<sup>a</sup>Membrane fractions from the indicated bacterial strains were assayed for the formation of [<sup>3</sup>H]GlcNAc-lipids in the presence of increasing amounts of Und-P, added as a dispersion in 1% CHAPS. Apparent kinetic parameters for Und-P were calculated from Michaelis-Menton plots (Fig. S5) using the linear regression algorithm in Sigma Plot ver. 12 (Systat Software, Inc. San Jose, CA). R squared values were 0.9953 for GAS GacI expressed in *E. coli*; 0.983 for GAS GacI co-expressed with GacJ in *E. coli*, 0.9976 for GacI in MGAS5005 and 0.931 for GacO in MGAS5005. The results are representative of at least two separate experiments. ND = Not Detected.

### GlcNAc-P-P-Und is required for initiation of polyrhamnose backbone biosynthesis

The GAS genome contains a close relative, *gacO*, of *E. coli wecA*, the tumicamycin-sensitive GlcNAc-phosphate transferase responsible for the synthesis of GlcNAc-P-P-Und in many bacteria (11,12). Significantly, GAC synthesis in GAS is inhibited by tunicamycin (5) and a close homolog of GacO, *S. mutans* RpgP, is required for rhamnopolysaccharide synthesis and can complement WecA deficiency in *E. coli* (13). Our analysis of UDP-[^3^H]GlcNAc incorporation in GAS and GBS membrane lipids, revealing the presence of two GlcNAc-lipids, GlcNAc-P-Und and GlcNAc-P-P-Und, prompted us to investigate the role of GlcNAc-P-P-Und in the initiation of GAC biosynthesis. First, we investigated the role of GacO in the synthesis of GlcNAc-P-P-Und. When membrane fractions from the WecA deficient *E. coli* strain (CLM37) expressing GacO, CLM37:GacO, were incubated with UDP-[^3^H]GlcNAc and Und-P (added as a dispersion in 1 % CHAPS), [^3^H]GlcNAc-P-P-Und was formed at an enzymatic rate that is similar to that found in the WecA over-expressor strain, PR4019 (Fig. 7A). CLM37 carrying an empty vector synthesizes no detectable GlcNAc-P-P-Und under these conditions (Fig. 7A).

**Figure 7.**
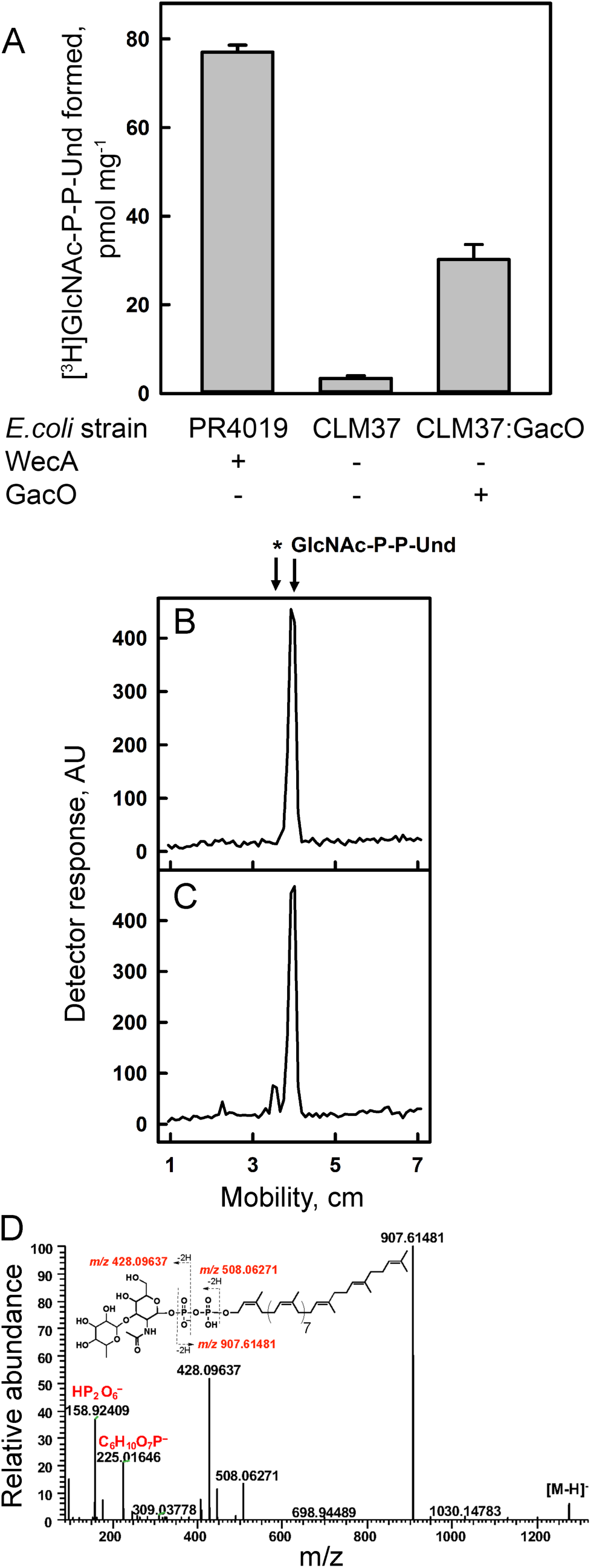
Analysis of GacO function in MGAS5005. (A) GacO catalyzes the synthesis of GlcNAc-P-P-Und in *E. coli* membranes. Reaction mixtures contained 50 mM Tris-HCl, pH 7.4, 5 mM 2-mercaptoethanol, 20 mM MgCl_2_, 0.5 % CHAPS, 20 μM Und-P (dispersed by ultrasonication in 1 % CHAPS), 5 μM UDP-GlcNAc (452 cpm/pmol) and *E. coli* membrane fraction from either PR4019, CLM37 or CLM37:GacO strains. Following incubation for 10 min at 30 °C, incorporation of [^3^H]GlcNAc into [^3^H]GlcNAc-P-P-Und was determined as described in Experimental procedures. Data are the average of three replicates ± standard deviation. (B and C) GlcNAc-P-P-Und can function as an acceptor substrate for rhamnosylation to form Rhamnosyl-GlcNAc-P-P-Und in 5005Δ*gacI* membranes. Incubation conditions were as described in the legend to Figure 3. (B) After 35 minutes incubation with UDP-[^3^H]GlcNAc, the reactions were analyzed for formation of [^3^H]GlcNAc lipids as described in Figure 3. (C) After 5 minutes incubation with UDP-[^3^H]GlcNAc, the reactions were incubated with 20 μM TDP-Rhamnose for an additional 30 min and analyzed for formation of [^3^H]GlcNAc lipids as described in Figure 3. The results are representative of three separate experiments. (D) ESI-MS/MS spectrum of rhamnosyl-GlcNAc-P-P-Und formed during chase with TDP-rhamnose from GlcNAc-P-P-Und synthesized *in situ* in membranes from *S. pyogenes* 5005Δ*gacI*.

To test whether GlcNAc-P-P-Und might function as a membrane anchor for the synthesis of the polyrhamnose chain of GAC, membrane fractions from MGAS5005 or the 5005Δ*gacI* mutant were pre-incubated with UDP-[^3^H]GlcNAc and ATP, to form [^3^H]GlcNAc-PP-Und *in situ*, and chased with non-radioactive TDP-rhamnose. The formation of the resultant [^3^H]GlcNAc-lipids in 5005Δ*gacI* was analyzed by TLC on silica gel G after 30 min incubation as shown in Fig. 7B and C. In the absence of TDP-rhamnose, 5005Δ*gacI* membranes produced only [^3^H]GlcNAc-P-P-Und (Fig. 7B), whereas in the presence of TDP-rhamnose two radioactive products, [^3^H]GlcNAc-P-P-Und and an additional product with slower mobility on TLC were observed (Fig 7C). MS analysis of lipid extracts from the *in vitro* reactions revealed the presence of a molecular ion [M-H]^-1^ M/z = 1274.76084 with the composition, C_69_H_114_O_16_NP_2_, consistent with the expected product, rhamnosyl-GlcNAc-P-P-Und (Figure 7D). Furthermore, the major fragment ions derived from M/z = 1274.76084 can be assigned to Rha-GlcNAc-P, Rha-GlcNAc-P-P and P-Und, supporting the proposed structural interpretation (Figure 7D and Table 4). Figure S6 shows the time-dependent accumulation of the new GlcNAc-lipid product. Significantly, parallel incubations with MGAS5005 showed that GlcNAc-P-Und was not glycosylated further. These results strongly support the possibility that GlcNAc-P-P-Und is an acceptor for rhamnosyl units in GAS and may function as the lipid anchor for the GAC polyrhamnose backbone synthesis.

**Table 4.**
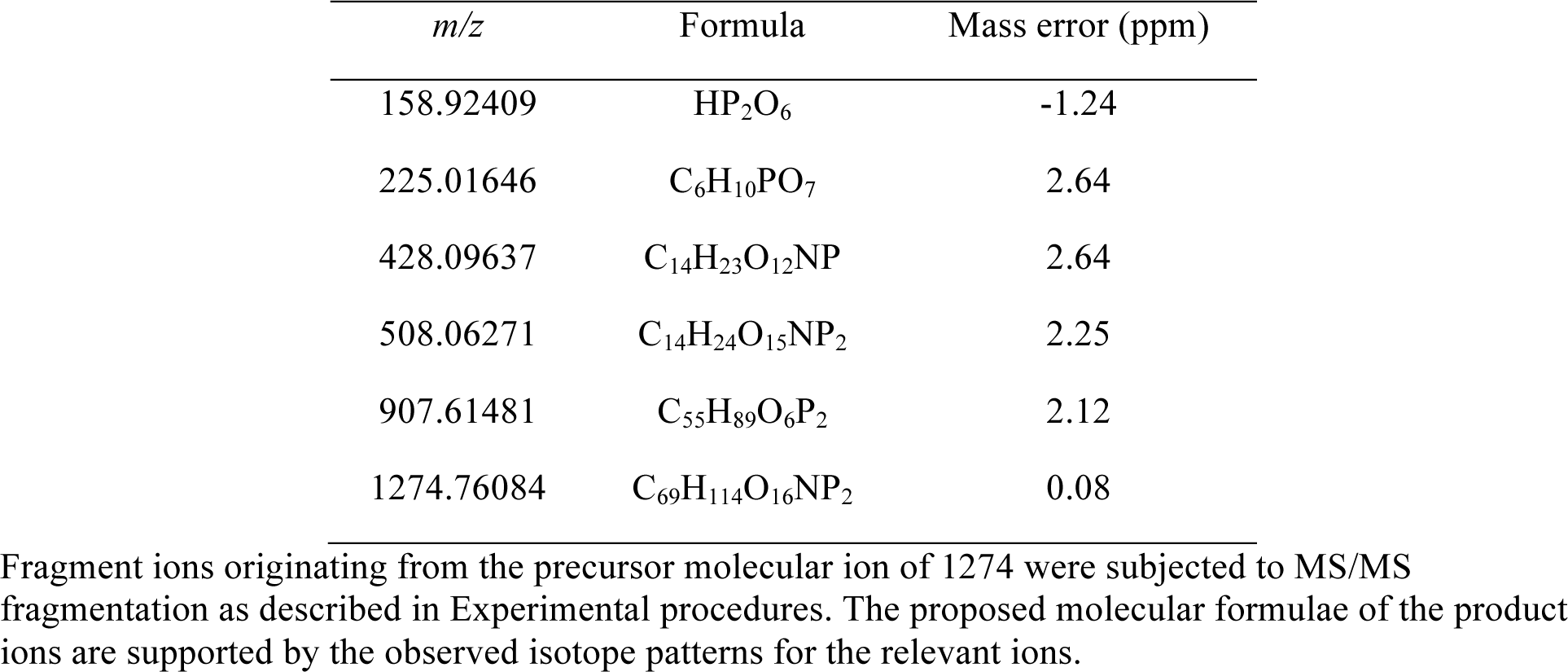
The major fragments of the precursor ion at *m/z* 1274.7612.

### The GacL mutant displays increased sensitivity to peptidoglycan amidases

The Gram-positive cell wall protects interior structures, plasma membrane and peptidoglycan, from host defense peptides and hydrolytic and antimicrobial enzymes. It has been shown that the GacI mutant which is GlcNAc-deficient is more sensitive to LL-37-induced killing (5). To test the hypothesis that the loss of GlcNAc decorations in GAC alters cell wall permeability, we investigated the sensitivity of 5005Δ*gacL* to peptidoglycan amidases: PlyC (27), PlyPy (28) and CbpD (29). When 5005Δ*gacL* cells were grown in increasing concentrations of CbpD (Fig. 8A), PlyPy (Fig. 8B) or PlyC (Fig. 8C) cellular growth was dramatically inhibited compared to MGAS5005, indicating increased sensitivity to the presence of amidases.

**Figure 8.**
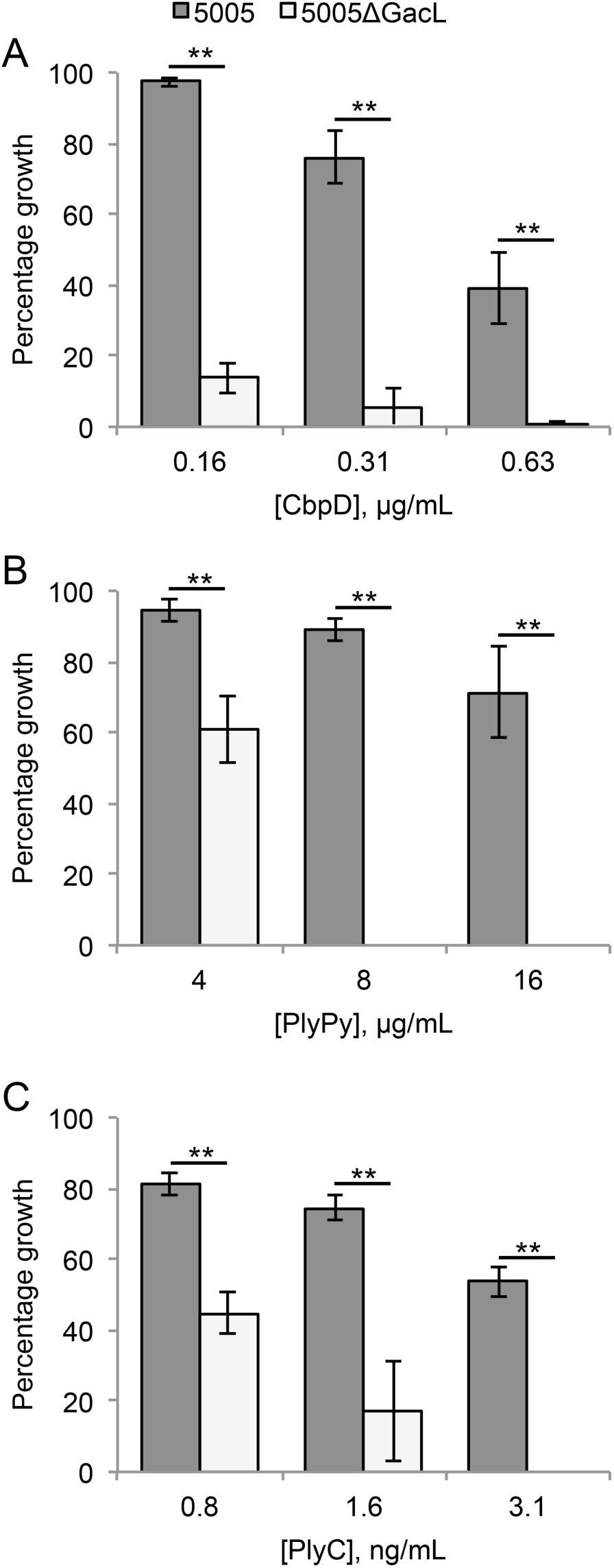
The absence of GacL increases sensitivity to cell wall amidases. Mid-exponential phase MGAS5005 and 5005Δ*gacL* were grown in the indicated concentrations of (A) CbpD, (B) PlyPy, and (C) PlyC. The change in growth is represented as a percentage of growth where no amidase was present. Data are the average of three replicates ± standard deviation. The asterisk indicate statistically different values (** p < 0.01) as determined by the Student’s t-test.

**Figure 9.**
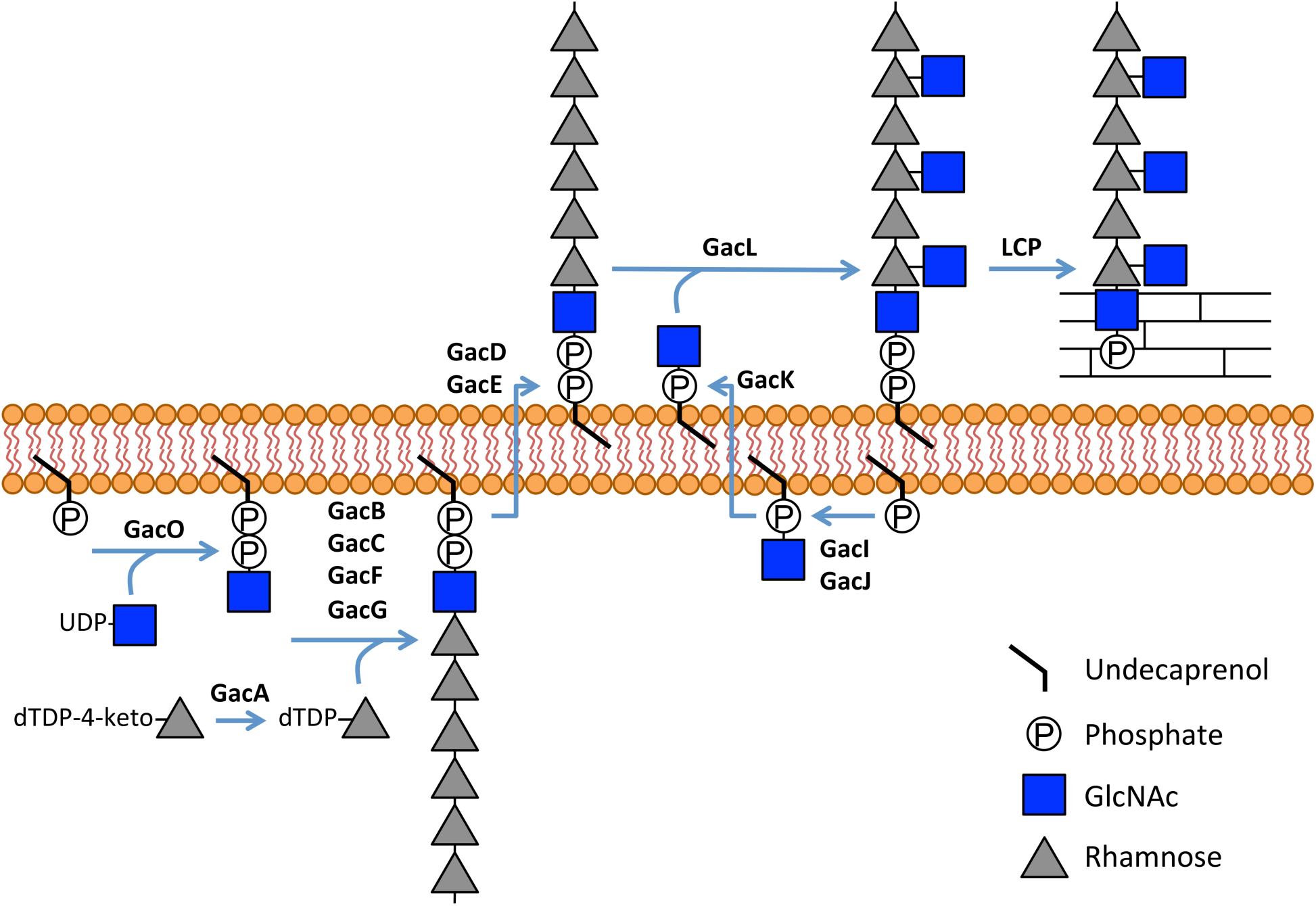
Schematic diagram of GAC biosynthesis. GAC is anchored to peptidoglycan presumably via phosphodiester bond. GAC biosynthesis is initiated on the inner leaflet of the plasma membrane where GacO produces GlcNAc-P-P-Und which serves as a membrane-anchored acceptor for polyrhamnose synthesis catalyzed by the GacB, GacC, GacF and GacG rhamnosyltransferases. Following polymerization polyrhamnose is transferred to the outer leaflet of the membrane presumably by the GacD/ GacE ABC transporter. Also in the inner leaflet of the membrane, GacI aided by GacJ produces GlcNAc-P-Und which then diffuses across the plasma membrane to the outer leaflet aided by GacK. Subsequently, GacL transfers GlcNAc to polyrhamnose using GlcNAc-P-Und as glycosyl donor. Lastly, protein members of LytR-CpsA-Psr phosphotransferase family presumably attach GAC to peptidoglycan. Several details of this biosynthetic scheme are still speculative and further research will be required to definitively confirm this hypothetical pathway, but the overall organization is consistent with other isoprenol-mediated capsular polysaccharide pathways.

## Discussion

In almost all Gram-positive bacteria, cell wall-attached glycopolymers are critical for cell envelope integrity and their depletion is lethal (30). Most streptococcal species including two human pathogens GAS and GBS do not synthesize WTA and instead produce Rha-containing glycopolymers as functional homologs of WTA (1). GAC is the major cell wall component of GAS and plays important roles in bacterial physiology and pathogenesis. The polyrhamnose core of GAC is modified with GlcNAc in an approximately 2:1 ratio (3,7) of rhamnose to GlcNAc. Collectively, the results of our study suggest a molecular mechanism of GAC biosynthesis in which rhamnan polymer is assembled at the cytoplasmic face of the plasma membrane, translocated to the cell surface and modified by GlcNAc on the outer side of the membrane as illustrated in Figure 9. We report that a lipid carrier, GlcNAc-P-P-Und, synthesized by GAS GacO, is a potential acceptor for initiation of rhamnan backbone biosynthesis. We speculate that the next step of rhamnan biosynthesis involves the action of the GacB, GacC, GacG and GacF glycosyltransferases. The lipid-anchored polyrhamnose is then translocated across the membrane by the ABC transporter encoded by *gacD* and *gacE*. This hypothesis is supported by studies of rhamnan biosynthesis in *S. mutans* (13,31,32).

Biosynthesis of the rhamnan-backbone of GAC is likely essential for GAS viability because the GacA enzyme involved in dTDP-rhamnose biosynthesis (14), and GacB and GacC glycosyltranferases are indispensable in GAS (5). In contrast, deletion of genes required for GlcNAc attachment to the rhamnan backbone does not affect GAS viability (5). This observation supports our hypothesis that biosynthesis of polyrhamnose and its translocation to the cell surface occur separately from the pathway involved in polyrhamnose modification with GlcNAc. In bacterial and eukaryotic glycoconjugate assembly systems, structural modifications that are introduced late in the biosynthesis typically involve polyprenyl monosphosphoryl donors (33). In our study, we found that GlcNAc modification of rhamnan requires GlcNAc-P-Und synthesis. Previously, GlcNAc-P-Und was isolated from various *Bacilli* membranes (12,23). However the enzyme required for GlcNAc-P-Und biosynthesis was not identified and the biological function of this lipid remained unclear. Our *in vitro* analysis of UDP-GlcNAc incorporation into GlcNAc-lipids by GAS membranes showed that GacI was required for the biosynthesis of GlcNAc-P-Und. Moreover CHAPS-soluble, affinity-purified GacI protein catalyzed the transfer of GlcNAc from UDPGlcNAc to Und-P yielding GlcNac-P-Und in an *in vitro* reaction system.

Interestingly, we found that, in contrast to *B. cereus*, GAS and GBS membranes incubated with UDP-[^3^H]GlcNAc synthesized primarily [^3^H]GlcNAc-P-Und. Since GacI and GacO utilize a common pool of Und-P, this phenomenon is presumably due to a much higher apparent affinity of GacI for Und-P compared to GacO. Significantly, we observed a measurable increase in [^3^H]GlcNAc incorporation into [^3^H]GlcNAc-P-P-Und in the GacI deletion mutant. This observation further confirmed that the relative synthetic rates of GlcNAc-P-Und and GlcNAc-P-P-Und biosynthesis are determined largely by relative affinity for Und-P. Furthermore, we found that the formation of GlcNAc-P-Und and GlcNAc-PP-Und in GAS membranes was significantly stimulated by ATP. Since GacI activity is not stimulated directly by ATP, it is likely that the effect of ATP is due to increased formation of Und-P, *in situ*, via undecaprenol kinase activity. In *S. mutans,* membrane-associated undecaprenol kinase catalyzes the ATP-dependent phosphorylation of undecaprenol to Und-P (26). It is likely that the homolog of this enzyme encoded by M5005_Spy_0389 is responsible for Und-P biosynthesis in GAS.

GacI is predicted by the HHpred server to have structural homology to polyisoprenylglycosyltransferase GtrB from *Synechocystis* which is a homolog of *S. flexneri* GtrB (34). GtrB homologs in bacteria and eukaryotes belong to the GT-A superfamily of glycosyltransferases and are responsible for the transfer of glucose from UDP-glucose to the bactoprenol carrier in the cytoplasm yielding the Und-P-glucose precursor in bacteria and dolichol phosphate-glucose precursor in eukaryotes (35,36). Our BLAST search using the GacI sequence as query found homologs of this gene in many bacterial species, with the broadest diversity observed in the phylum Firmicutes (supplemental file 1). To determine evolutionary relationships between the GacI homologs, these sequences were analyzed using CLANS (CLuster ANalysis of Sequences) (37) (Fig. 10). It is noteworthy that the *B. cereus* GacI has two homologs clustered in two different groups. Our bioinformatics analysis shows that these *gacI* homologs are located in different gene clusters encoding proteins involved in polysaccharide biosynthesis and transport. Moreover, the GAS GacI subgrouped with one *B. cereus* GacI. The GBS GacI and *E. faecalis* GacI homologs are located in the cluster well-separated from the GAS GacI subgroup. Thus, this observation points to possible horizontal gene transfer between GAS and *Bacillus*. Analysis of UDP-GlcNAc incorporation into GlcNAc-lipids in a GBS *gacI* knock-out strain confirmed the function of this gene in the biosynthesis of GlcNAc-P-Und in GBS. Additionally, we demonstrated that the *E. faecalis* GacI homolog (EpaI) functions in the biosynthesis of GlcNAc-P-Und. Significantly, the presence of GacI homologs in GBS and *E. faecalis* matches the reported occurrence of GlcNAc residues in the cell wall polysaccharides of these bacteria (38-41).

**Figure 10.**
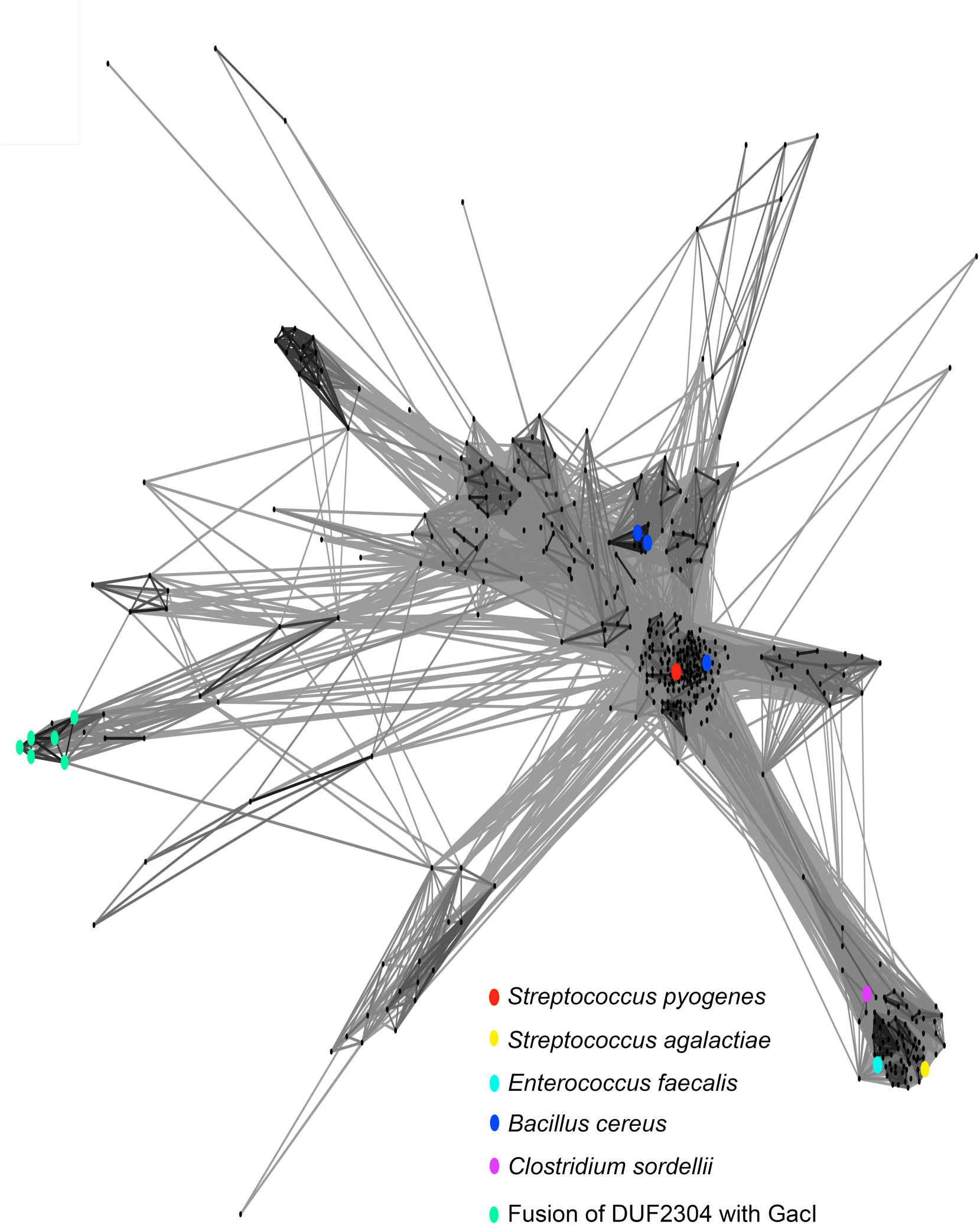
Sequence relationship of GacI family of proteins. Homology between GacI homologs is graphically displayed using CLANS analysis (37). Dots correspond to individual protein sequences selected as described in Experimental procedures and provided as Supplementary File. Selected homologs of *S. pyogenes* are highlighted by colored dots.

GacJ, a small membrane protein with three transmembrane α-helices, is required for GlcNAc side-chain attachment to polyrhamnose (5). GacJ belongs to the DUF2304 family of proteins according to the Pfam protein family database (42). In *Geobacter* sp., *Desulfuromonas* sp., *Desulfurivibrio alkaliphilus, Desulfuromonas soudanensis* and *Desulfurispirillum indicum*, DUF2304 homologous domains are fused with *gacI* homologs (Fig. 10 and supplemental file 1). In *M. tuberculosis* a GacJ homolog, Rv3632, is co-transcribed with the gene encoding N-acetyl galactosaminyl-phosphate-undecaprenol synthase, PpgS, and is found to stimulate PpgS activity (43). Consistent with this finding we showed that GacI and GacJ form a complex and co-expression of GacI with GacJ significantly enhanced GacI catalytic activity. Further work is under way to characterize the mechanism of GacJ action on GacI activity.

In many bacterial systems modification of glycoconjugates involves the export of polyprenyl monosphosphoryl-glycose molecule to the periplasm providing the glycosyl donor for the modifying reaction catalyzed by integral membrane glycosyltransferase that possesses a GT-C fold (33). Bioinformatics searches with the HHpred program against available protein structures identified the GT-C superfamily of glycosyltransferases as the closest structural homologs of GacL. Moreover, the phenotypes of the GacL mutant: absence of the GlcNAc side-chains in GAC and accumulation of GlcNAc-P-Und in the membrane is consistent with GacL functioning in the transfer of GlcNAc from GlcNAc-P-Und to polyrhamnose yielding GlcNAc-modified polysaccharide.

We suggest that the Wzx flippase, GacK, may function to transport GlcNAc-P-Und to the extracellular space for utilization as GlcNAc donor in the GlcNAc modification of polyrhamnose. This hypothesis is based on the GlcNAc-deficient phenotype of the GacK mutant (5). However, further research will be required to confirm this hypothesis. The last step of GAC biosynthesis is probably similar to the last step of WTA biosynthesis: it involves attachment of GAC to certain MurNAc residues in peptidoglycan via a phosphate ester linkage (46). It is likely catalyzed by members of LytRCpsA-Psr (LCP) phosphotransferase family encoded by M5005_Spy_1099 and M5005_Spy_1474.

The identification of the mechanism of GlcNAc attachment to polyrhamnose raises the question of how this modification functions biologically in bacteria. It has been previously found that the Δ*gacI* mutant is hypersusceptible to human antimicrobial peptide LL-37 and the antimicrobial action of factors released by thrombin-activated platelets, suggesting a role for GlcNAc modification in protecting the plasma membrane from antimicrobial agents (5). Our study identified the importance of GlcNAc for protection of GAS peptidoglycan from amidase-induced lysis. In *E. faecalis* the GacI homolog, EpaI, is involved in biosynthesis of a cell wall-attached polysaccharide (24), however the polysaccharide structure of the mutant has not been elucidated. The Δ*epaI* mutant was defective in conjugative transfer of a plasmid and resistance of bacteria to detergent and bile salts (24).

In conclusion, our study provides a platform for elucidation of novel pathways of glycopolymer biosynthesis in other bacterial pathogens including important drug-resistant bacteria such as *E. faecalis* and *Clostridium sordellii* that possess GacI and GacJ homologs (Fig. 10 and supplemental file 1). Since the enzymes involved in the biosynthesis of cell wall attached glycopolymers are promising targets for novel antimicrobials and the glycopolymers represent important features for diagnostics and vaccine targets, our data may provide opportunities for developing novel therapeutics against antibiotic-resistant bacterial pathogens.

## Experimental procedures

*Bacterial strains and growth conditions:* All plasmids, strains and primers used in this study are listed in Tables S2 and S3 in the supplemental material. The strains used in this study were GAS M1-serotype strain MGAS5005 (16), *Bacillus cereus* (ATCC 14579), *Streptococcus agalactiae* COH1 (Group B *Streptococcus* or GBS), *E. coli* CLM37 (47), *E. coli* PR4019 (12), *E. coli* DH5α and *E. coli* Rosetta (DE3). GAS and GBS cultures were grown in Todd-Hewitt broth (BD) supplemented with 0.2% yeast extract (THY), or on THY agar plates at 37 °C. *E. coli* and *B. cereus* strains were grown in Luria-Bertani (LB) medium or on LB agar plates at 37 °C. When required, antibiotics were included at the following concentrations: ampicillin at 100 μg ml^-1^ for *E. coli*; streptomycin at 100 μg ml^-1^ for *E. coli*; erythromycin at 500 μg ml^-1^ for *E. coli* and 1 μg ml^-1^ for GAS and GBS; chloramphenicol at 10 μg ml^-1^ for *E. coli* and 5 μg ml^-1^ for GAS; spectinomycin at 200 μg ml^-1^ for *E. coli* and 100 μg ml^-1^ for GAS and GBS.

*DNA techniques:* Plasmid DNA was isolated from *E. coli* by commercial kits (Qiagen) according to the manufacturer’s instructions and used to transform *E. coli*, GAS and GBS strains. Plasmids were transformed into GAS and GBS by electroporation as described previously (48). Chromosomal DNA was purified from GAS and GBS as described in (49). To construct single-base substitutions or deletion mutations, we used the QuikChange^®^ II XL Site-Directed Mutagenesis Kit (Stratagene) according to the manufacturer’s protocol. Constructs containing mutations were identified by sequence analysis. All constructs were confirmed by sequencing analysis (Eurofins MWG Operon).

*Construction of the* gacI *deletion mutant in GAS:* For construction of strain 5005Δ*gacI*, MGAS5005 chromosomal DNA was used as a template for amplification of two DNA fragments using two primers pairs: GacIm-BamHI-f/GacIdel-r and GacIdel-f/GacIm-XhoI-r (Table S3). Primer GacIdel-f is complementary to primer GacIdel-r. The two gel-purified PCR products containing complementary ends were mixed and amplified using a PCR overlap method (50) with primer pair GacIm-BamHI-f/GacIm-XhoI-r to create the deletion of *gacI*. The PCR product was digested with BamHI and XhoI and ligated into BamHI/SalI-digested temperature-sensitive shuttle vector pJRS233 (51). The plasmid was designated pJRS233Δ*gacI* (Table S2). The resulting plasmid was transformed into MGAS5005, and erythromycin resistant colonies were selected on THY agar plates at 30 °C. Integration was performed by growth of transformants at 37 °C with erythromycin selection. Excision of the integrated plasmid was performed by serial passages in THY media at 30 °C and parallel screening for erythromycin-sensitive colonies. Mutants were verified by PCR sequencing of the loci.

*Construction of the* gacL *deletion mutant in GAS:* To create BglII and XhoI cloning sites in the polylinker region of pUC19, site-directed mutagenesis of the plasmid, using two primer pairs, pUC19-XhoI-f/pUC19-XhoI-r and pUC19-BglII-f/pUC19-BglII-f, was carried out. The plasmid was designated pUC19BX. The nonpolar *aadA* spectinomycin resistance cassette was amplified from pLR16T (Table S2) using primers Spec-SalI-f/Spec-BamH-r (Table S3), digested with SalI/BamHI, and ligated into SalI/BamHI-digested pUC19BX to yield pUC19BXspec. MGAS5005 chromosomal DNA was used as a template for amplification of two DNA fragments using two primers pairs: gacLup-BglII-f/gacLup-SalI-r and gacLdown-BamHI-f/gacLdown-XhoI-r (Table S3). The first PCR product was digested with BglII/SalI and ligated into BglII/SalI-digested pUC19BXspec. The resultant plasmid, pUC19BXspecL1, was digested with BamHI/XhoI and used for ligation with the second PCR product that was digested with BamHI/XhoI. The resultant plasmid, pUC19BXspecL2, was digested with BglII and XhoI to obtain a DNA fragment containing *aadA* flanked with the *gacL* upstream and downstream regions. The DNA fragment was ligated into pHY304 vector (52) and digested with BamHI/XhoI to yield pHY304Δ*gacL*. The resulting plasmid was transformed into MGAS5005, and erythromycin resistant colonies were selected on THY agar plates at 30 °C. The mutants were isolated as describe above. 5005Δ*gacL* mutants were screened for sensitivity to spectinomycin and verified by PCR sequencing of the loci.

*Complementation of the 5005Δ*gacL *mutant with* gacL *(5005Δ*gacL gacL^*+*^): To construct the plasmid for complementation of the 5005Δ*gacL* mutant, MGAS5005 chromosomal DNA was used as a template for amplification of a wild-type copy of *gacL* using the primer pair GacLXhoI-f and GacL-BglII-r (Table S3). The PCR products were digested with XhoI and BglII and cloned in pBBL740 (Table S2) previously digested with the respective enzymes. The integrational plasmid pBBL740 does not have a replication origin that is functional in GAS, so the plasmid can be maintained only by integrating into the GAS chromosome through homologous recombination. The resultant plasmid pGacL was transformed into 5005Δ*gacL* by electroporation and transformants were selected on agar plates containing chloramphenicol. Several chloramphenicol resistant colonies were selected and *gacL* integration into the chromosome was confirmed by sequencing a PCR fragment.

*Construction of the* gacI *(SAN_1536) deletion mutant in GBS:* GBS COH1 chromosomal DNA was used as a template for amplification of two DNA fragments using two primers pairs: gbsIup-BglII-f/gbsIup-SalI-r and gbsId-BamHI-f/gbsId-XhoI-r (Table S3). The plasmid for *gacI* (SAN_1536) knock-out was constructed using the same strategy described for *gacL* deletion (see above). The resultant plasmid, pHY304GBSΔI, was transformed into GBS COH1, and erythromycin resistant colonies were selected on THY agar plates at 30 °C. The mutants were isolated as described above. GBS COH1 Δ*gacI* mutants were screened for sensitivity to spectinomycin and verified by PCR sequencing of the loci.

*Construction of the plasmids for* E. coli *expression of GacI, GacJ and GacO:* To create a vector for expression of GacI from GAS, the gene was amplified from MGAS5005 chromosomal DNA using the primer pair GacINcoI-f and GacI-XhoI-r (Table S3). The PCR product was digested with NcoI and XhoI, and ligated into NcoII/XhoI-digested pRSF-NT vector (Table S3). The resultant plasmid, pGacI, contained *gacI* fused at the N-terminus with a His-tag followed by a TEV protease recognition site. To create a vector for expression of GacI and GacJ, the bicistronic DNA fragment was amplified from MGAS5005 chromosomal DNA using the primer pair GacI-NcoI-f and GacJXhoI-r (Table S3). The vector was constructed as described above. The plasmid was designated pGacIJ. pET21_NESG plasmid for expression of the GacI homolog, EpaI, from *Enterococcus faecalis* was obtained from the DNASU repository (53). The construct was confirmed by sequencing analysis. The plasmids were transferred into competent *E. coli* Rosetta (DE3) (Novagen) using the manufacturer’s protocol.

To create a vector for expression of GacO, the gene was amplified from MGAS5005 chromosomal DNA using the primer pair GacOXba-f and GacO-HindIII-r (Table S3). The PCR product was digested with XbaI and HindIII, and ligated into XbaI/HindIII-digested pBAD33 vector. The resultant plasmid, pGacO, was transferred into competent *E. coli* CLM37 strain that has a deletion of the *wecA* gene.

*Construction of the plasmids for* E. coli *expression of PlyPy, CbpD and PlyC:* To create a vector for expression of CbpD amidase (29), the gene was amplified from MGAS5005 chromosomal DNA using the primer pair 28-NcoI-f and 28-stop-r (Table S3). The PCR product was digested with NcoI and XhoI, and ligated into NcoI/XhoI-digested pRSF-NT vector. The resultant plasmid, pCbpD, contained *cbpD* fused at the N-terminus with a His-tag followed by a TEV protease recognition site. To create a vector for expression of PlyPy amidase (28), the gene was amplified from MGAS5005 chromosomal DNA using the primer pair PlyPy-NcoI-f and PlyPy-XhoI-r. The PCR product was digested with NcoI and XhoI, and ligated into NcoI/XhoI-digested pET-21d vector. The resultant plasmid, pPlyPy contained the gene fused at the C terminus with a His-tag.

To create a vector for expression of PlyC amidase (54), a DNA fragment spanning bicistronic operon which encodes *plyCA* and *plyCB* (27) was synthesized by ThermoFisher Scientific. The plasmid was digested with NcoI and XhoI, and ligated into NcoI/XhoI-digested pET21d vector. The resultant plasmid, pPlyC contained *plyCA* followed by *plyCB* fused at the C terminus with a His-tag.

*Expression and purification of GacI, EpaI and GacI/GacJ complex:* For expression of GacI, EpaI and GacI/GacJ complex, *E. coli* Rosetta (DE3) cells carrying the respective plasmid were grown to an OD_600_ of 0.4-0.6 and induced with 1 mM IPTG at 18 °C for approximately 16 hours. The cells were lysed in 20 mM Tris-HCl pH 7.5, 300 mM NaCl with two passes through a EmulsiFlex-C5 microfluidizer cell disrupter (Avestin, Inc., Ottawa Ontario,CA). The lysate was centrifuged at 7000 x g for 30 minutes, 4°C. The supernatant was centrifuged at 30,000 x g for 60 minutes, 4°C to isolate the membrane fraction.

To isolate GacI/GacJ complexes the membrane proteins were solubilized in 2.5% CHAPS, 20 mM Tris-HCl pH 7.5, 300 mM NaCl for 60 minutes, rotating at room temperature. Insoluble material was removed by centrifugation at 30,000 x g for 60 minutes, 4°C. Solubilized GacI/GacJ were purified by Ni-NTA chromatography with washes of 2.5% CHAPS, 20 mM Tris-HCl pH 7.5, 300 mM NaCl and 2.5% CHAPS, 20 mM Tris-HCl pH7.5, 300 mM NaCl, 10 mM imidazole, and elution with 2.5% CHAPS, 20 mM Tris-HCl pH 7.5, 300 mM NaCl, 250 mM imidazole. The GacI/GacJ complex was further purified by size exclusion chromatography on a Superdex 200 (GE Biosciences) column in 2.5% CHAPS, 20 mM HEPES pH 7.5, 100 mM NaCl, with monitoring for protein elution at 280 nm.

*Expression of GacO:* For expression of GacO, *E. coli* CLM37 cells carrying the pertinent plasmid were grown to an OD_600_ of 0.8 and induced with 13 mM L-arabinose at 25 °C for approximately 3 hours. The cells were lysed in 20 mM Tris-HCl pH 7.5, 300 mM NaCl with two passes through a microfluidizer cell disrupter. The lysate was centrifuged at 1000 x g for 15 minutes, 4 °C. The supernatant was centrifuged at 40,000 x g for 60 minutes, 4 °C to isolate the membrane fraction.

*Expression and purification of PlyPy, CbpD and PlyC;* For expression and purification of PlyPy, CbpD and PlyC, *E. coli* Rosetta (DE3) cells carrying the respective plasmids were grown to an OD_600_ of 0.4-0.6 and induced at 18 °C with 1 mM IPTG for approximately 16 hours. The cells were lysed in 20 mM Tris-HCl pH 7.5, 300 mM NaCl with two passes through a microfluidizer cell disrupter. The soluble fraction was purified by Ni-NTA chromatography with washes of 20 mM Tris-HCl pH 7.5, 300 mM NaCl and 20 mM Tris-HCl pH7.5, 300 mM NaCl, 10mM imidazole, and elution with 20 mM Tris-HCl pH 7.5, 300 mM NaCl, 250 mM imidazole. The PlyPy and CbpD eluate was further purified by size exclusion chromatography on a Superdex 200 column in 20 mM HEPES pH 7.5, 100 mM NaCl for PlyPy, or 20 mM MOPS pH 6.5, 100 mM NaCl for CbpD. PlyC was further purified by anion exchange chromatography on a MonoQ 5/50 GL (GE Biosciences) column in 10 mM sodium phosphate pH 6.0, and a 20 column volume elution gradient of 0-500 mM NaCl.

*Isolation of GAS and GBS membranes:* Bacteria were grown at 37 °C to an OD_600_ of 0.8. To obtain GAS cell membranes, cell pellet was re-suspended in phosphate-buffered saline (PBS) and incubated 1 h with PlyC lysin as described in (55). To obtain GBS membranes, the cell pellet was re-suspended in acetate buffer (50 mM CH_3_COONa, 10 mM CaCl, 50 mM NaCl, pH 5) and incubated with mutanolysin (200 U/ml, Sigma–Aldrich) for 2 h at 37 °C. After hydrolytic enzyme treatment the bacterial suspension was sonicated using a Fisher Scientific™ Model 505 Sonic Dismembrator, 15 times ×15 s. After centrifugation at 8,000 *g* for 10 min at 4 °C, the supernatant was collected and centrifuged at 40,000 *g* for 60 min. The pellet was collected as the membrane fraction.

*Mass-spectrometry analysis of GacI and GacJ:* LC-MS/MS for proteomic analysis was performed using an LTQ-Orbitrap mass spectrometer (Thermo Fisher Scientific) coupled with an Eksigent Nanoflex cHiPLC system (Eksigent) through a nanoelectrospray ionization source. The LC-MS/MS data were subjected to database searches for protein identification using Proteome Discoverer software V. 1.3 (Thermo Fisher Scientific) with a local MASCOT search engine.

*Dot-blot analysis:* Bacterial cells (1 ml) from exponential phase cultures (OD_600_=0.8) was centrifuged, washed with PBS, resuspended in 100 μl of PBS and incubated with 1 μl PlyC (1.5 mg/ml) for 1 h at 37°C. After centrifugation at 16,000 *g* for 2 min, 5 μl of the supernatant was spotted on a nitrocellulose membrane. The membrane was blocked 1 h with 7% skim milk in PBS with 0.1% Tween 20, and incubated overnight with an anti-GAC antibody diluted 1:5,000 (ab9191; Abcam). Bound antibodies were detected with a peroxidase-conjugated goat anti-rabbit immunoglobulin G antibody, and the Amersham ECL (enhanced chemiluminescence) Western blotting system.

*Binding of succinylated wheat germ agglutinin (sWGA) to bacteria:* MGAS5005 and 5005Δ*gacL* cells were collected during mid-exponential phase (OD_600_=0.6), washed three times with BSA-saline solution (0.5% BSA, 0.15 M NaCl), resuspended in BSA-saline solution and incubated for 30 minute at 37 °C with mixing. GlcNAc specific fluorescein-sWGA was added to final concentrations of 0, 12.5, 25, 50, 100 μg/ml. After 1 h of incubation at 37 °C with mixing, the cells were centrifuged, washed twice, and resuspended in BSA-saline. Bound fluorescein-sWGA was quantified in a fluorimeter SpectraMax M5 (Molecular Devices) using an excitation of 544 nm and emission of 590 nm.

*Isolation of cell wall from GAS:* MGAS5005 and 5005Δ*gacL* cell wall was prepared from exponential phase cultures (OD_600_=0.8) by the SDS-boiling procedure as described for *Streptococcus pneumoniae* (56). Purified cell wall samples were lyophilized and used for composition analysis.

*Carbohydrate composition analysis:* Carbohydrate composition analysis was performed at the Complex Carbohydrate Research Center (Athens, GA) by combined gas chromatography/mass spectrometry (GC/MS) of the per-O-trimethylsilyl (TMS) derivatives of the monosaccharide methyl glycosides produced from the sample by acidic methanolysis as described previously by (57).

*Assay of lytic activity of PlyPy, CbpD and PlyC:* Aliquots of MGAS5005 and 5005Δ*gacL* frozen stocks [prepared as described in (58)] were inoculated into THY medium 1:20. To a 96 well plate, 200 μL aliquots of culture were dispensed in duplicate, per condition. The plates were grown at 37 °C without aeration for 2 hours. An assay plate was set up containing PlyPy (8 μg mL^-1^, 16 μg mL^-1^, 32 μg mL^-1^), CbpD (0.31 μg mL^-1^, 0.63 μg mL^-1^, 1.26 μg mL^-1^) or PlyC (1.6 ng mL^-1^, 3.1 ng mL^-1^, 6.2 ng mL^-1^) in a volume of 100 μL. To the assay plate, 100 μL of culture was added to each well giving final concentrations of: PlyPy (4 μg mL^-1^, 8 μg mL^-1^, 16 μg mL^-1^), CbpD (0.16 μg mL^-1^, 0.31 μg mL^-1^, 0.63 μg mL^-1^), or PlyC (0.8 ng mL^-1^, 1.6 ng mL-^1^, 3.1 ng mL^-1^). After mixing, the absorbance at 600 nm was measured for each well (t_0_) and the plate was incubated at 37 °C for 1 h. The absorbance at 600 nm was then measured again (t_1_). The percentage difference (t_1_-t_0_) in growth relative to 0 mM of enzyme/antimicrobial, for each concentration was calculated (i.e. 0 mM PlyPy = 100%).

*Extraction and characterization of bacterial glycolipids:* Two liters of bacterial cells from exponential phase cultures (OD_600_=0.8) were recovered by sedimentation at 10,000 x g for 30 min, and washed with ice-cold PBS 2x. Cells were resuspended in 50 ml PBS, sensitized by incubation with 10 ng mL^-1^ PlyC for 1 h at 37 °C and stirred vigorously with two volumes of CH_3_OH and one volume of CHCl_3_ for 30 min at room temperature. The mixture was divided into 5 mL aliquots in 12 mL glass centrifuge tubes. Insoluble material was removed by centrifugation at 200 *g* and the organic extract was transferred to a separatory funnel. The insoluble residue was further extracted with 1 mL of CHCl_3_/CH_3_OH (2:1) per tube, two times, and the extracts were combined with the previous organic phase. The organic extract was supplemented with CHCl_3_ and 0.9% NaCl to give a final composition of CHCl_3_/CH_3_OH/0.9% NaCl (3:2:1), mixed vigorously, and allowed to stand in the cold until phase separation was achieved. The organic phase was drained off into a second separatory funnel and the organic layer was washed with 1/3 volume of CHCl_3_/CH_3_OH/0.9% saline (3:48:47), two times. The aqueous layers were discarded. The organic extract was dried on a vacuum rotary evaporator, dissolved in a small volume of CHCl_3_/CH_3_OH (2:1) and transferred to a 12.5 x 100 mm screw cap glass tube (with Teflon lined cap). The organic extract was dried under a stream of nitrogen gas and the glycerolipids were destroyed by deacylation in 0.1 M KOH in toluene/CH_3_OH (1:3) at 0 °C, 60 min. Following deacylation, the reactions were neutralized with acetic acid, diluted with two volumes of CHCl_3_,1 volume of CHCl_3_/CH_3_OH (2:1) and 1/5 volume of 0.9% NaCl/10 mM EDTA. The two-phase mixture was mixed vigorously and centrifuged to separate the phases. The organic phase was dried under nitrogen, spotted on a 20 x 20 cm sheet of Silica Gel G and developed in CHCl_3_/CH_3_OH/H_2_O/NH_4_OH (65:25:4:1). Bacterial lipids were visualized by staining with iodine vapors and pertinent spots were scraped from the thin layer plate, eluted from the silica gel with CHCl_3_/CH_3_OH (2:1) and reserved for further analysis.

*Mass spectrometry analysis of a phospholipid isolated from 5005Δ*gacL: A phospholipid accumulated by 5005Δ*gacL* was purified as described above using preparative TLC in silica gel. The compound was analyzed by LC-MS using a Q-exactive mass spectrometer and an Ultimate 3000 ultra high performance liquid chromatography system (Thermo Fisher Scientific, San Jose, CA) on a Kinetex C18 reversed-phase column (2.6 mm × 100 mm, 2.1 μm, Phenomenex, USA). Two solvents were used for gradient elution: (A) acetonitrile/water (2:3, v/v), (B) isopropanol/acetonitrile (9:1, v/v). Both A and B contained 10 mM ammonium formate and 0.1% formic acid. The column temperature was maintained at 40 °C, and the flow rate was set to 0.25 ml/min. Mass spectrometric detection was performed by electrospray ionization in negative ionization mode with source voltage maintained at 4.0 kV. The capillary temperature, sheath gas flow and auxiliary gas flow were set at 330 °C, 35 arb and 12 arb, respectively. Full-scan MS spectra (mass range m/z 400 to 1500) were acquired with resolution R = 70,000 and AGC target 5e5. MS/MS fragmentation was performed using high-energy C-trap dissociation with resolution R = 35,000 and AGC target 1e6. The normalized collision energy was set at 30.

*Assay for incorporation of GlcNAc into polyisoprenol-linked glycolipid intermediates:* Reaction mixtures for measuring the incorporation of [^3^H]GlcNAc into lipids contained 50 mM Tris, pH 7.4, 0.25 M sucrose, 20 mM MgCl_2_, 5 mM β-mercaptoethanol, 5 μM UDP-[6-^3^H]GlcNAc [(100-2000 cpm/pmol) American Radiolabelled Chemicals] and bacterial membrane suspension (50-250 μg bacterial membrane protein) in a total volume of 10 to 100 μL. In some experiments, 1 mM ATP was included, as indicated. Following incubation at 30 °C, the enzymatic reactions were terminated by the addition of 40 volumes of CHCl_3_/CH_3_OH (2:1), thoroughly mixed and incubated for 5 min at room temperature. Insoluble material was sedimented at 200 *g* and the organic extract was transferred to a 12x100 mm glass tube. The residue was re-extracted with 1 ml CHCl_3_/CH_3_OH (2:1), two times, and the organic extracts were combined. The pooled organic extracts were freed of unincorporated radioactivity by sequential partitioning with 1/5 volume of 0.9% saline/10 mM EDTA and then with 1/3 volume of CHCl_3_/CH_3_OH/0.9% saline (3:48:47) 3 times, discarding the aqueous phase each time. The washed organic phases were dried under nitrogen and re-dissolved in CHCl_3_/CH_3_OH (2:1). A carefully measured aliquot was removed and analyzed for radioactivity by liquid scintillation spectrometry after drying. The remainder of the sample was analyzed by thin layer chromatography on Silica Gel G, developed in CHCl_3_/CH_3_OH/H_2_O/NH_4_OH (65:25:4:1), using a BioScan AR2000 radiochromatoscanner. Incorporation of [^3^H]GlcNAc into individual [^3^H]GlcNAc-lipids was calculated using the peak integration values obtained from the BioScan.

*Degradation of GlcNAc-lipids by mild acid and mild alkaline treatment:* Bacterial lipids were subjected to mild acid hydrolysis (50% isopropanol, 0.1 M HCl, 50 °C, 1 h) and mild alkaline methanolysis [0.1 M KOH in toluene/CH_3_OH (1:3), 0 °C, 1 h]. Following incubation, the reactions were neutralized with either 1 M Tris or concentrated acetic acid, diluted with CHCl_3_, CH_3_OH and 0.9% NaCl to give a final composition of 3:2:1, respectively and partitioned as described above. The organic phases were dried under nitrogen and either quantified for radioactivity or analyzed by thin layer chromatography and detected by iodine staining.

*Phenolysis of GlcNAc-P-Und:* Purified [^3^H]GlcNAc-P-Und was dried under nitrogen in a 12x100 mm conical screw-cap tube and heated to 68 °C in 0.2 mL 50% aqueous phenol as described by Murazumi et al. (22). Following phenolysis, 0.1 ml water was added and the samples were thoroughly mixed. The aqueous and phenolic layers were separated, dried and analyzed for radioactivity by scintillation spectrometry.

*Analytical methods:* Membrane protein concentrations were determined using the BCA protein assay kit (Pierce Chemical Co.) employing the method of Ruiz et al. (44). Radioactivity was quantified by liquid scintillation spectrometry on a Packard Tri-Carb Liquid Scintillation Spectrometer using Econosafe Complete Counting Cocktail (Research Products International, Inc., Elk Grove IL).

*Phylogenetic analysis of GacI homologs:* Sequences of GacI homologs were retrieved from the non-redundant database using BLAST (59). One thousand sequences were downloaded and were manually curated to remove sequences with shorter that 90% sequence length overlap. The redundancy was reduced using CD-HIT with 0.98 cut off (60). Sequences were manually curated to correct misannotated translation start codons. The pair-wise similarities were analyzed and visualized using CLANS with an E-value cut off 1 e^-^80 (37,61,62).

*Bioinformatics analysis:* The TOPCONS (http://topcons.net/) (63) web server was employed to predict trans-membrane regions of GacI, GacJ and GacL. Homology detection and structure prediction were performed by the HHpred server (https://toolkit.tuebingen.mpg.de/#/tools/hhpred) (64).

*Statistical analysis:* Unless otherwise indicated, statistical analysis was carried out from at least three independent experiments. Quantitative data was analyzed using the paired Student’s t-test. A P value equal to or less that 0.05 is considered statistically significant.

## Acknowledgements

This work was supported by NIH grants R21AI113253 from the National Institute of Allergy and Infectious Diseases (to NK) and R01GM102129 (to Dr. Charles J. Waechter and JSR), 1S10OD021753 to AJM and by the Center of Biomedical Research Excellence (COBRE) Pilot Grant (to KVK, NK and JSR) supported by NIH grant P30GM110787 from the National Institute of General Medical Sciences. NMvS is supported by VIDI grant 91713303 from the Netherlands Organization for Scientific Research (NWO).

Carbohydrate composition analysis at the Complex Carbohydrate Research Center was supported by the Chemical Sciences, Geosciences and Biosciences Division, Office of Basic Energy Sciences, U.S. Department of Energy grant (DE-FG02-93ER20097) to Parastoo Azadi.

The authors thank Dr. Charles J. Waechter for encouragement and helpful discussions.

The funders had no role in study design, data collection and interpretation, or the decision to submit the work for publication.

## Author contributions

JSR, RJE, HZ, AJM, NMvS, KVK and NK designed the experiments. JSR, RJE and NK performed biochemical experiments. PD, JC, HZ and AJM performed MS analysis. NK constructed plasmids and isolated mutants. JSR, RJE, KVK and NK wrote the manuscript. All authors reviewed the results and approved the final version of the manuscript.

